# Grains, trade and war in the multimodal transmission of Rice yellow mottle virus: an historical and phylogeographical retrospective

**DOI:** 10.1101/2024.08.20.608750

**Authors:** Innocent Ndikumana, Geoffrey Onaga, Agnès Pinel-Galzi, Pauline Rocu, Judith Hubert, Hassan Karakacha Wéré, Antony Adego, Mariam Nyongesa Wéré, Nils Poulicard, Maxime Hebrard, Simon Dellicour, Philippe Lemey, Erik Gilbert, Marie-José Dugué, François Chevenet, Paul Bastide, Stéphane Guindon, Denis Fargette, Eugénie Hébrard

## Abstract

Rice yellow mottle virus (RYMV) is a major pathogen of rice in Africa. RYMV has a narrow host range limited to rice and a few related poaceae species. We explore the links between the spread of RYMV in East Africa and rice history since the second half of the 19^th^ century. The phylogeography of RYMV in East Africa was reconstructed from coat protein gene sequences (ORF4) of 335 isolates sampled over two million square kilometers between 1966 and 2020. Dispersal patterns obtained from ORF2a and ORF2b, and full-length sequences converged to the same scenario. The following imprints of rice cultivation on RYMV epidemiology were unveiled. RYMV emerged in the middle of the 19^th^ century in the Eastern Arc Mountains where slash-and-burn rice cultivation was practiced. Several spillovers from wild hosts to cultivated rice occurred. RYMV was then rapidly introduced into the adjacent large rice growing Kilombero valley. Harvested seeds are contaminated by debris of virus infected plants that subsist after threshing and winnowing. Long-distance dispersal of RYMV is consistent (i) with rice introduction along the caravan routes from the Indian Ocean Coast to Lake Victoria in the second half of the 19^th^ century, (ii) seed movement from East Africa to West Africa at the end of the 19^th^ century, from Lake Victoria to the north of Ethiopia in the second half of the 20^th^ century and to Madagascar at the end of the 20^th^ century, (iii) and, unexpectedly, with rice transport at the end of the First World War as a troop staple food from the Kilombero valley towards the South of Lake Malawi. Overall, RYMV dispersal was associated to a broad range of human activities, some unsuspected. Consequently, RYMV has a wide dispersal capacity, its dispersal metrics estimated from phylogeographic reconstructions are similar to those of highly mobile zoonotic viruses.

**Author summary:** Rice yellow mottle virus (RYMV) poses a major threat to rice production in Africa. We explored through a multidisciplinary approach the links between the history of rice in East Africa since the second half of the 19^th^ century and the spread of RYMV. The results illuminate the causes of RYMV diffusion. We show the role of long-distance caravan trade, the impact of the First World War and the consequences of seed exchange in the dispersal of RYMV. The paradoxical role of seeds in the spread of RYMV - which is vector transmitted and not seed transmitted – is explained in the light of rice biology and agronomy. Overall, this study reveals the wide range of transmission ways, some unexpected, in the dispersal of plant viruses. It also highlights the role of human transmission of pathogens, even vector-borne, and sheds light on the risk of transmission of RYMV and of other plant viruses from Africa to other continents.

## Introduction

Contact transmission is a major transmission pathway for plant viruses, yet it has largely been neglected [1]. Contact-transmitted viruses have particles that are stable outside the infected cells, remain infectious on contaminated surfaces, and reach high concentrations (titer) in infected plants. Owing to their virion stability and their high titer, Sobemoviruses, Tobamoviruses, Potexviruses and Tombusviruses are listed as candidates for contact and abiotic transmission [1]. The sobemovirus species differ from the three other genera by a narrow host range and by the mode of transmission, most often by beetles. Beetles are involved in the transmission of several genera [2].

Rice yellow mottle virus (RYMV) is a single-stranded RNA virus of the Sobemovirus genus [3]. RYMV is a major pathogen in all sub-Saharan African countries that grow rice [4]. Its host range is limited to the two cultivated rice species *Oryza sativa* and *O. glaberrima*, to the wild rice species and to a few related *Eragrostideae* species [5]. RYMV does not infect the seed embryo [6, 7, 8]. RYMV is transmitted by chewing insects, mainly beetles of the Chrysomelidae family, in a non-persistent mode and by leaf and root contact between infected and healthy plants [5]. RYMV reached high virus concentration within two to three weeks. It remains infectious in rice stubbles for months [5, 7], and after passage through the intestinal tract of mammals [9]. The spread of the disease was first attributed to vector transmission, although no clear link has been found between the size of the beetle populations and the incidence of the disease. Later, the role of mechanical transmission through leaf-to-leaf and root-to-root contact was established experimentally [10]. Contamination in the soil also occurred by contact of the roots of young seedlings with infected rice stubble present in the soil. This is referred to as soil transmission or soil contamination [1, 11, 12]. Soil contamination has several origins, infected rice residues buried in the soil after harvesting or after mammal consumption, and also by planting rice seeds contaminated by infected plant residues (see below). Due to the stability of RYMV [7], soil transmission persists months after the infected rice residues have been buried. RYMV is disseminated by transplanting young infected seedlings from nurseries to fields sometimes located kilometers away [13]. Altogether, contact and soil transmission accounts for on-site survival of the virus and its local spread.

Yet, the long-distance transmission of RYMV has not been elucidated. Without experimental evidence, it has been tentatively ascribed to movements of beetles during tornadoes, transmission by more proficient flyer insects such as grasshoppers [5], or transmission by seeds in specific circumstances [14, 15]. The last of the three hypotheses, which posited transmission with seeds, relied on the following observations. RYMV is not present in the rice embryo but is detected in the glumes, and in the outer envelopes of the grain [8]. Rice threshing and winnowing does not eliminate all the outer husks and all the leaf and stem residues [16]. Accordingly, infected plant debris remain in bags of rice seeds. We referred to them as contaminated seeds to make the distinction with infected seeds [4, 17]. Rice seeds are transported between villages and across regions for food consumption and for agronomical purposes [18]. Rice planting with contaminated seeds as well as consumption of contaminated rice seeds by mammals (including humans) result in burying infected rice residues in the soil. Rice planted in a contaminated soil is infected through mechanical contact of the roots of the young seedlings with the infected residues. Based on these observations, it was hypothesized that RYMV was transported with bags of contaminated rice seeds to new areas, potentially far away from its sources [4, 17]. Accordingly, seed transport would be the primary mode, possibly the exclusive mode, of long distance dissemination of RYMV.

Nevertheless, long-distance transmission of plant viruses, be it by vectors or with seeds, has been difficult to establish [19, 20], and this is true for RYMV as well. None of the hypotheses put forward for long-distance transmission of RYMV has been experimentally validated. Phylogeography offers new perspectives through the reconstruction of the spatiotemporal pattern of spread [21, 22]. It unveils the underlying processes that shape the dynamics of spread, including rare and past events, and leads to hypotheses on the modes of transmission and the impact of host biology and history. RYMV is a measurably evolving population [23], and the phylogeographic reconstructions are not affected by the time dependent rate phenomenon [24]. The relaxed random walk (RRW) model in continuous phylogeography allows for variation in dispersal rates across branches of the phylogeny [22]. This model provides flexibility to accommodate the different means of transmission when reconstructing the phylogenetic dispersal history of the virus.

In this study, the phylogeography of RYMV in East Africa was reconstructed from a comprehensive heterochronous sequence dataset. Given that RYMV has a narrow host range limited to rice and a few related species, it was expected that its dissemination has been shaped by rice history. Subsequently, we paid special attention to the history of rice in East Africa since the beginning of the 19^th^ century. RYMV originates in East Africa but unlike in Madagascar and in West Africa, rice cultivation had remained localized until recently [25, 26, 27]. Rice had been cultivated along the Indian Ocean Coast for centuries but was confined to a few scattered coastal sites [26]. The sole documented case of large plantation at the beginning of the 19^th^ century is the Kilombero valley, upstream of the mouth of the coastal river Rufiji, 300 km from the Indian Ocean [28]. By the end of the 19^th^ century, rice supplies from the Rufiji flood plains had emerged as a sufficiently important source for that area to acquire the nickname “Little Calcutta” [29, 30]. In the middle of the 19th century, rice was introduced around the great lakes of East Africa (Lake Victoria, Lake Tanganyika and Lake Malawi) [31]. Rice cultivation became generalized at the end of the 20^th^ century. Assessing the virus movements between these far apart rice-producing regions helps to identify the main modes of dissemination.

Dispersal statistics are increasingly used to characterise and quantify the spread of viruses [32, 33, 34]. We assessed the dispersal capacity of RYMV and compare it to a range of zoonotic viruses [33]. Altogether, this study suggests that the transport of contaminated rice seed was decisive in the long-distance dissemination of RYMV, not only within East Africa, but also in the introduction of the virus from East Africa to West Africa and to Madagascar. It points the most probable routes and modes of dissemination and unveils the links with rice history. It further shows the likely role of a broad spectrum of human activities in RYMV dispersal, some of them unsuspected. This explains why RYMV exhibits extensive dissemination capabilities, similar to those of highly mobile zoonotic viruses.

## Results

### Temporal signal

BETS analyses indicate that the increase in sample size of heterochronous sequences lead to a stronger temporal signal in West Africa (from 261 to 335 sequences) and in East Africa (from 240 to 335 sequences) (Table 1). The temporal signal was more pronounced in the set of West African sequences (reflected by a higher Bayes factor). The temporal signal of the two datasets was also tested using cluster randomization tests of the sampling dates. There was no overlap of the credibility intervals of the substitution rate calculated with sampled dates and those with randomized dates (data not shown). However, the differences obtained were more pronounced with the WA335 dataset. The square of the correlation coefficient of the regression sampling date vs. distance to the root obtained with TreeTime was 0.07 with the WA335 dataset vs. 0.01 with the EA335 dataset. Altogether, the temporal signal was more pronounced in the WA335 dataset than in the EA335 dataset. The stronger temporal signal of the West African dataset probably results from a relatively uniform distribution of the sampling dates over the last thirty years whereas that of East Africa isolates was concentrated over the last 15 years. Short sampling periods are prone to unpurged transitory deleterious polymorphism leading to higher and biased substitution rates [35]. Consistently, the weak temporal signal of the EA335 dataset leads to a biased high substitution rate and correlatively to an inconsistent, obviously too recent, age of the most recent common ancestor of the sampled sequences (tMRCA). Given the nested position of the West African lineage within the East African lineages (see below), the tMRCA in East Africa is expected to be substantially older than that in West Africa.

**Table 1.** Temporal signals in the datasets assessed by BETS.

| Datasets | Log marginal likelihood* |  | Bayes factor |
| --- | --- | --- | --- |
|  | isochronous | dated |  |
| West-Africa |  |  |  |
| WA261 | -10495 | -10409 | 86 |
| WA335 | -13546 | -13379 | 167 |
| East-Africa |  |  |  |
| EA240 | -11294 | -11246 | 48 |
| EA335 | -14634 | -14539 | 95 |
| Madagascar |  |  |  |
| Mg94 | -3701 | -3671 | 30 |

\*estimated under a strict molecular clock model.

As the temporal signal from the East African corpus was not sufficient alone to allow a valid reconstruction of the spatio-temporal dispersion of the RYMV in East Africa, a prior was set on the substitution rate. The substitution rate estimates of the three datasets (WA261, WA335 and Mg94) were similar (Table 1). The substitution rate of the Mg94 dataset indicates that a reliable estimate was obtained from a dataset with a limited temporal signal after the integration of another data source, in this instance epidemiological information derived from field surveys. We selected as a prior the substitution rate of the WA335 dataset as its standard deviation was the lowest. Accordingly, for the EA335 phylogeographic reconstructions, a prior was set on the mean rate under the uncorrelated log-normal relaxed molecular clock (the “ucld.mean” parameter), a normal distribution with a mean of 1.24 10^-3^ substitutions per site per year and standard deviation of 1.16 10^-4^. A prior was also set on the standard deviation of the uncorrelated log-normal relaxed clock (the “ucld.stdev” parameter), a normal distribution with a mean of 1.09 x 10^-3^ substitution per site per year and standard deviation of 2.87 x 10^-4^.

### Phylogeny of RYMV in East-Africa

The phylogeny of RYMV in East Africa reconstructed from 335 coat protein gene sequences is made of three major lineages named, respectively, S4, S5 and S6 lineages (Fig. 1). The phylogenetic group containing the S4 and S5 lineages does not have a strong bootstrap support. Several strains are distinguished within the S4 and S6 lineages. The S4 lineage includes strains S4lm, S4lv, S4ug and S4et named according to their geographical distribution (lm, Lake Malawi; lv, Lake Victoria; ug, Uganda; et, Ethiopia). The S6 lineage includes strains S6a, S6b, and S7 but was not supported by a strong bootstrap value. The S5 lineage consists of one strain only.

**Figure 1.**
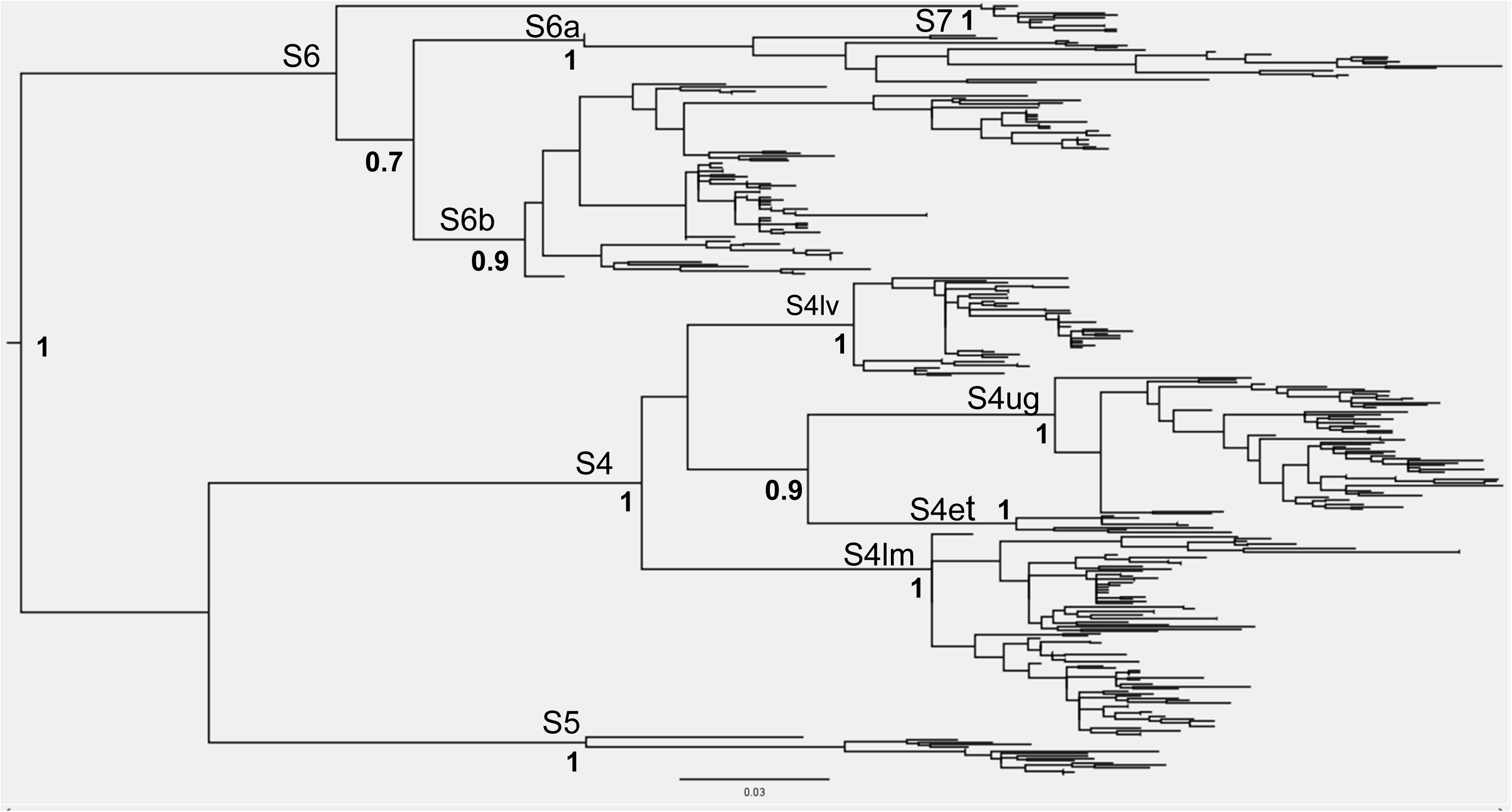
Phylogeny of RYMV in East Africa reconstructed from the coat protein gene sequences of 335 isolates through the maximum likelihood method under a HKY85 model and mid-point rooting. The names of the lineages and of the strains are given. The bootstrap support (> 0.70) is indicated at the node of the lineages and of the strains.

The phylogeny of RYMV was also reconstructed from the full-length genomic sequences (with recombination regions removed; see Materials and Methods) of 50 isolates representative of the geographic and genetic diversity of RYMV in East Africa (Fig. 2). The same phylogenetic pattern was observed at the lineage and strain levels, except that the S4lv and S4lm strains belong to the same phylogenetic group in the reconstruction from the full-length sequences but not from the coat protein gene sequences. Very few insertion-deletion events occurred in coding regions. The basal insertion-deletion pattern at positions 19 and 60 in the ORF4 in the S4, S5 and S6 lineages was observed earlier [36]. An additional deletion occurred in the isolate Tz319 of the S5 strain at codon 432 in the ORF2a. It is the only insertion-deletion event found at a terminal phylogenetic position (Fig. 2). All the insertion-deletion polymorphisms originated in isolates of the Eastern Arc Mountains.

**Figure 2.**
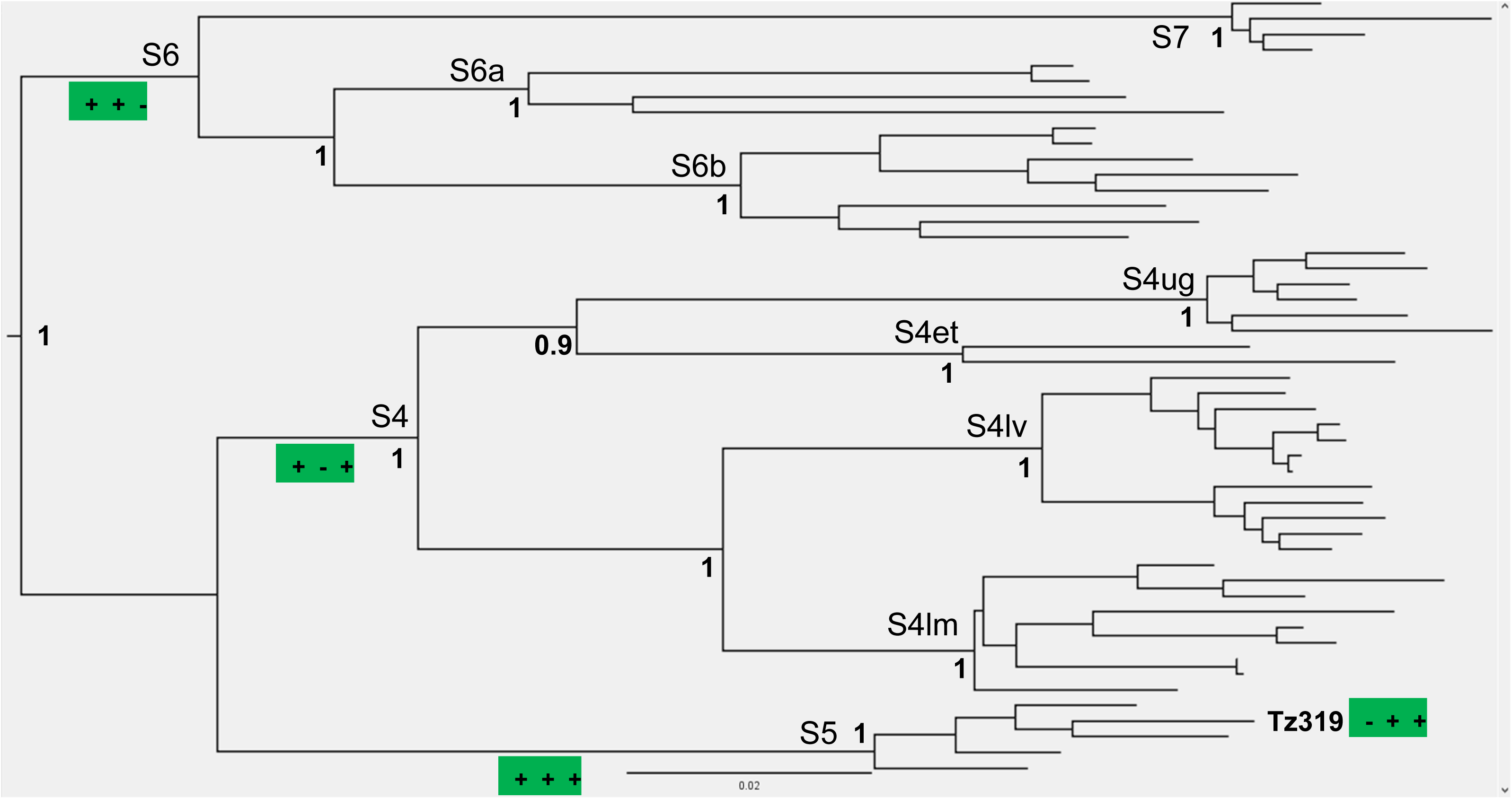
Phylogeny of RYMV in East Africa reconstructed from the full-length genome (with recombination regions removed; see Materials and Methods) of 50 isolates representative of the geographic and genetic diversity through a maximum likelihood method under a HKY85 model and mid-point rooting. The names of the lineages and of the strains are given. The insertion-deletion polymorphism of the lineages and of the isolate Tz319 are framed in green as followed: “+”, presence; “-”, absence of an amino acid at position 432 of the ORF2a, 19 and 60 of the ORF4 successively.

### Phylogeography from 335 coat protein gene sequences

The spatio-temporal dispersal of RYMV lineages in East Africa was reconstructed from the capsid protein gene sequences of 335 isolates (Figs. 3, 4, 5). The following periods are distinguished.

1. In the middle of the 19^th^ century, RYMV emerged around the Great Ruaha Escarpment in the Udzungwa mountains of the Eastern Arc Mountains north of the Kilombero Valley (Fig. 4);
2. Within a few decades, RYMV dispersed to the adjacent Kilombero Valley (Fig. 4);
3. Between the end of the 19^th^ century and the beginning of the 20^th^ century, RYMV reached the south of Lake Victoria 400 km northward of the Kilombero Valley, the first long range dispersal event (I) (lineage S4), and the Morogoro region 100 km eastward of the Kilombero Valley (lineage S6) (Fig. 4);
4. In the following decades, RYMV circulated at the east of Lake Victoria (Fig. 5);
5. A second long range dispersal event (II) originated at the beginning of the 20th century in the Kilombero valley to reach the south of Lake Malawi, 900km southward, 80 years later (Figs. 3. and 5);
6. A third long range dispersal event (III) took place in the second half of the 20th century. It originated at the east of Lake Victoria and reached northern Ethiopia, 1400 km northward, 50 years later (Figs. 3 and 5);
7. During the last decades of the 20th century, RYMV dispersed eastward and northward from the Kilombero Valley and the Morogoro region, and southward and westward from Lake Victoria (Fig. 4). Recently, the virus became widespread in all rice-growing regions with an increase in movements between adjacent localities (Figs. 3 and 5).

**Figure 3.**
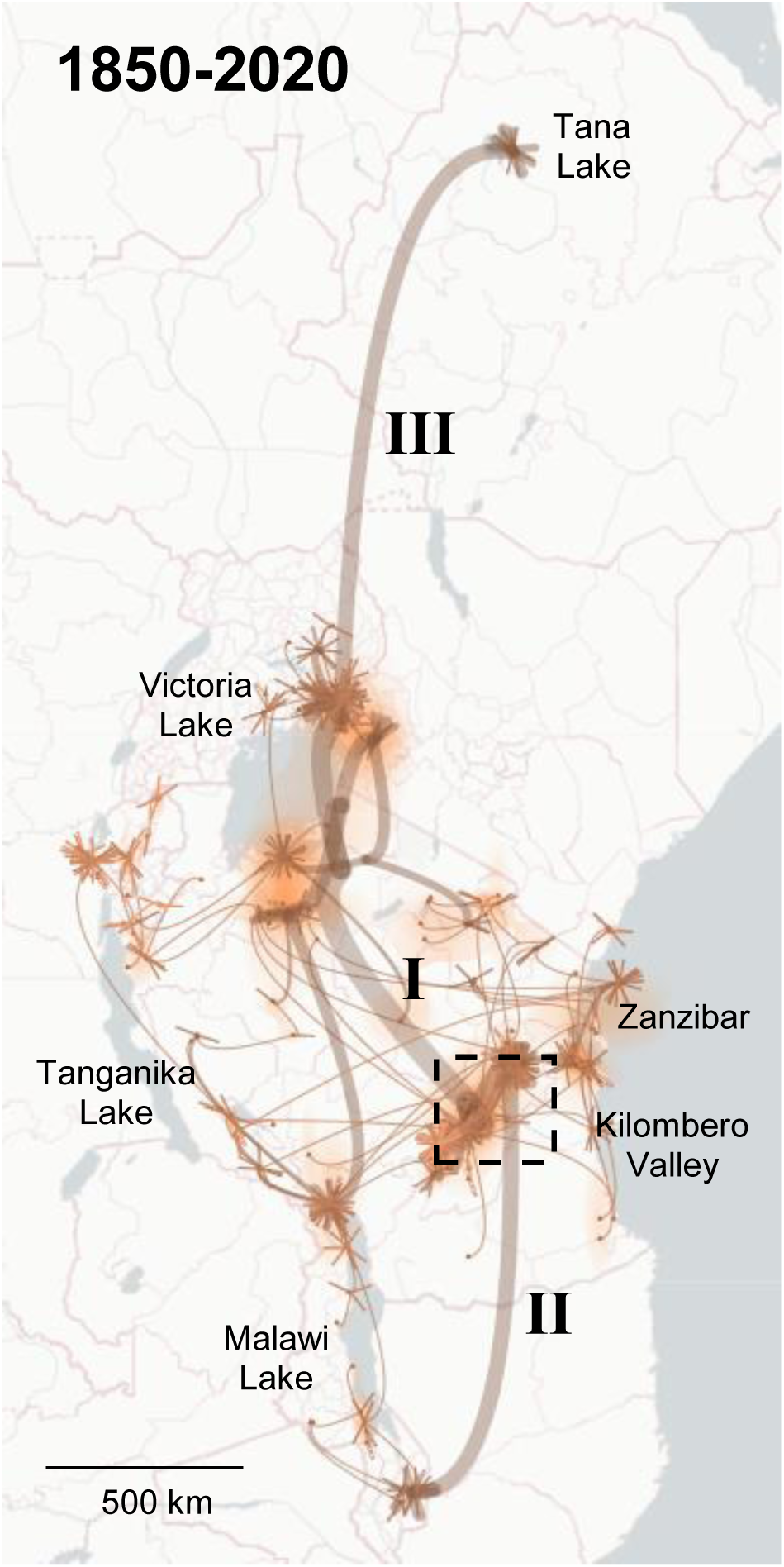
Continuous phylogeographic reconstruction of the spatiotemporal dispersal of RYMV lineages in East Africa from 1850 to 2020 based on the coat protein gene sequences of 355 isolates collected from 1966 to 2019. The maximum clade credibility (MCC) tree and the 95% highest posterior density (HPD) regions reflecting the uncertainty related to the phylogeographic inference were mapped. The phylogeographic scenario is displayed with temporal gradients linked to line thickness, curvature, opacity, and color. These gradients match the depth of the nodes in the phylogenetic tree, ranging from ancient to recent: path thickness ranges from wide to narrow, curvature varies from high to low, opacity shifts from light to strong, and color transitions from light to deep brown. The three long range dispersal events were indicated (I, II, III) and the RYMV emergence zone was figured by a dotted frame.

**Figure 4.**
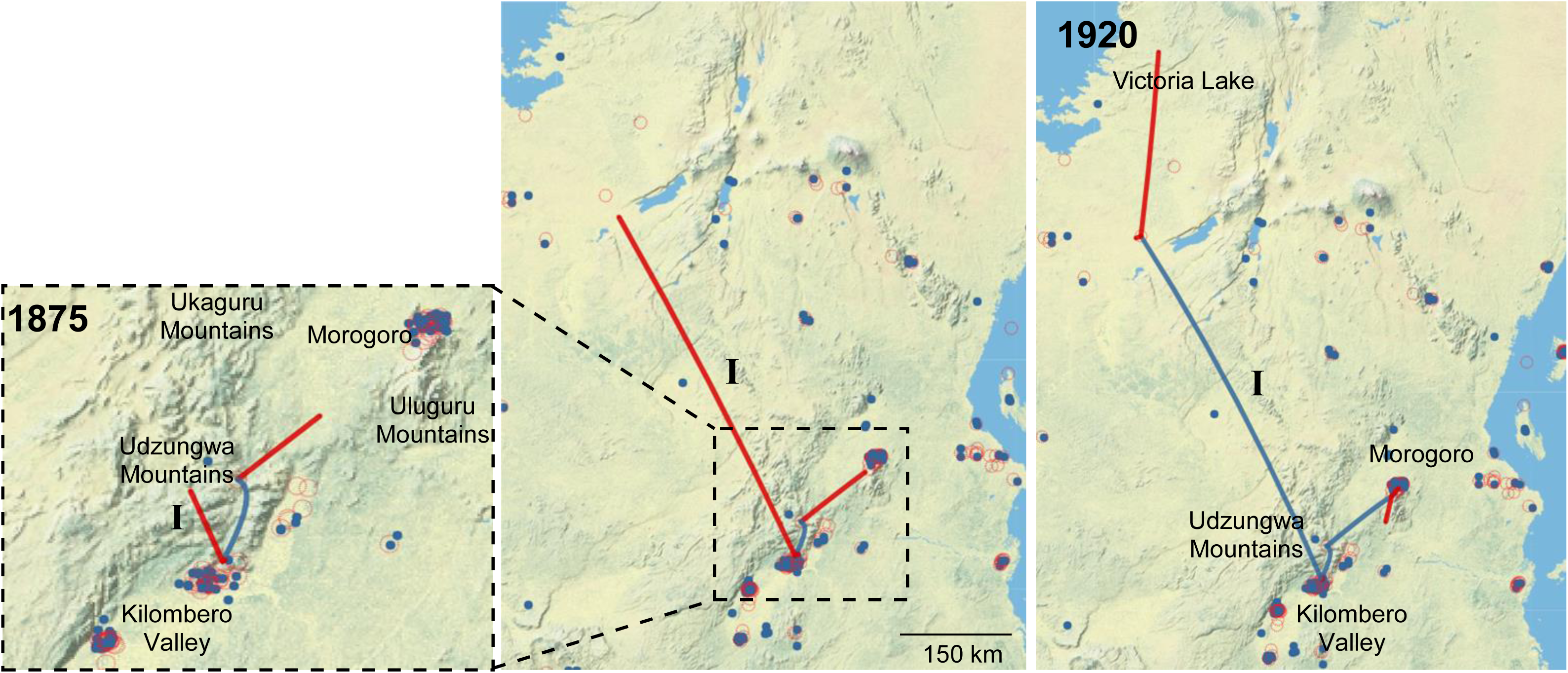
Visualization of the continuous phylogeographic reconstruction of RYMV lineages in East Africa from 1875 to 1920. Transitions associated with the category ‘parent node selected/child node not selected’ are highlighted in red to distinguish the ongoing migrations from the completed ones highlighted in blue and associated to the category ‘parent node selected/child node selected’. The location of the ancestral states (open red circle) is indicated alongside the location of the sampled isolates (full blue circles). The first long range dispersal event was indicated (I) and the RYMV emergence zone was visualized by a dotted frame.

**Figure 5.**
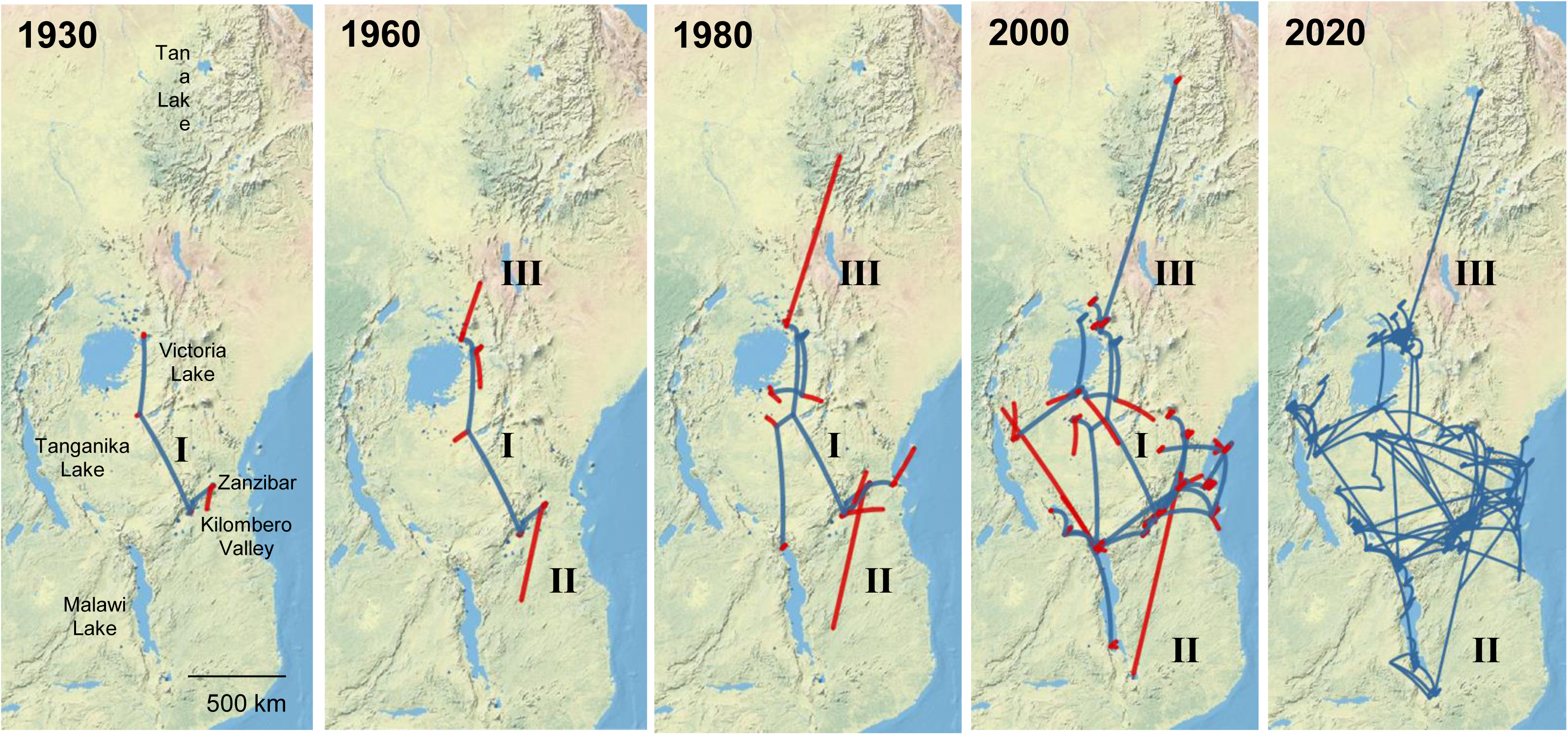
Visualization of the continuous phylogeographic reconstruction of RYMV lineages in East Africa from 1930 to 2020 at stages that capture the major dispersal events. Transitions associated with the category ‘parent node selected/child node not selected’ are highlighted in red to distinguish the ongoing migrations from the completed ones highlighted in blue and associated to the category ‘parent node selected/child node selected’.

Altogether, the phylogeography of RYMV in East Africa reconstructed from the 335 coat protein gene sequences was characterized by these three long-distance dispersal events along a south-north axis from the middle of the 19^th^ century to the second half of the 20^th^ century. In the past decades, a regular short distance dispersal eastward and westward became predominant.

### Phylogeography from full-length sequences and ORFs of 50 representative isolates

The spatio-temporal dispersal of RYMV in East Africa was reconstructed from sequences of 50 isolates representative of the genetic and geographic diversity of the virus in East Africa: full-length sequences (4468 nt, and excluding recombinant regions), sequences of ORF2a (including protease and VPg; 459 codons), ORF2b (including polymerase; 540 codons) and ORF4 (capsid protein, 240 codons). The differences in nucleotide diversity and in the selection pressure among the ORFs are given in the Supplementary Table 5. No recombination events was detected within the ORF2a, ORF2b and ORF4.

To make up for the weak temporal signal of these small datasets (50 sequences), a temporal prior was applied to the root of each tree by using the tMRCA of the EA335 tree, following a normal distribution with a mean 170 years and a standard deviation of 25 years. Regardless the differences in sequence length, diversity and selection pressure between the ORFs (Supplementary Table 5), the phylogeographic patterns were similar (Fig. 6), and consistent with that reconstructed from the 335 sequences of the coat protein gene: (1) an emergence in the Eastern Arc Mountains, (2) a rapid spread to the adjacent large rice growing valley of Kilombero, (3-4) dispersal northward towards Lake Victoria, and eastward towards the Morogoro region, (5) spread towards the south of Lake Malawi, (6) spread towards northern Ethiopia, (7) generalized dissemination throughout East Africa.

**Figure 6.**
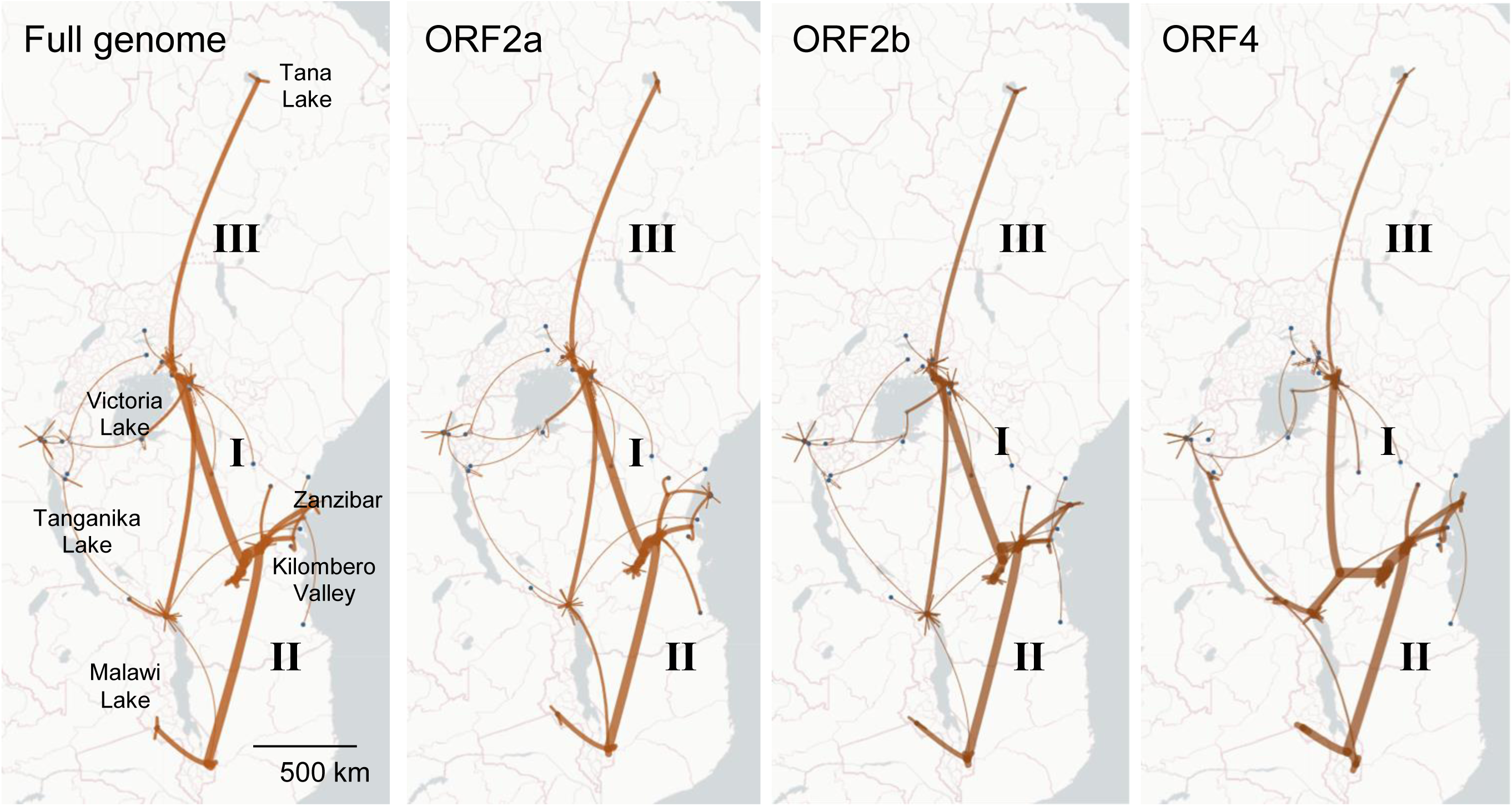
Continuous phylogeographic reconstructions of the spatiotemporal dispersal of RYMV lineages in East Africa from 1850 to 2020 based on the non-recombinant full-length sequences (a), ORF2a (b) and ORF2b (c) and ORF4 (d) of 50 representative isolates of the genetic and geographic diversity. The recombinant regions were removed from the full-length sequences (see Materials and Methods). The phylogeographic scenario is displayed with temporal gradients linked to line thickness, curvature, opacity, and color. These gradients match the depth of the nodes in the phylogenetic tree, ranging from ancient to recent: path thickness ranges from wide to narrow, curvature varies from high to low, opacity shifts from light to strong, and color transitions from light to deep brown.

### Phylogeography of the three main lineages

The initial diversification of RYMV in East Africa resulted in the emergence of the three lineages S4, S5 and S6 in the mid-19th century within or near the Kilombero valley. The phylogeography of the three lineages is markedly different (Figs. 7a, 7b). The S4 lineage had the greatest epidemic success. It was found south of Lake Victoria in the second half of the 19th century (S4lv), and then circulated around Lake Victoria (S4ug) before dispersing northward into Ethiopia (S4et), then southward to Malawi and finally westward to the Republic of Congo, Rwanda and Burundi (S4lm). The S6 lineage remained confined to the Kilombero valley and in the Morogoro region for several decades. In the last decades, it spread towards eastern Tanzania, south-west Kenya and to the islands of Zanzibar and Pemba. The S7 strain split early from the S6 lineage and was recovered recently in southern Malawi. The S5 lineage was found exclusively in the Kilombero valley. The migration curve (Fig. 7c) illustrates the differences in velocity among the lineages, with an early and rapid spread of the S4 lineage, a late but rapid spread of the S6 lineage (except the early spread of the S7 strain), and a lack of dispersal for S5. Interestingly, there was a slowdown in the dispersal rates the past decades within most strains of the S4 and S6 lineages.

**Figure 7.**
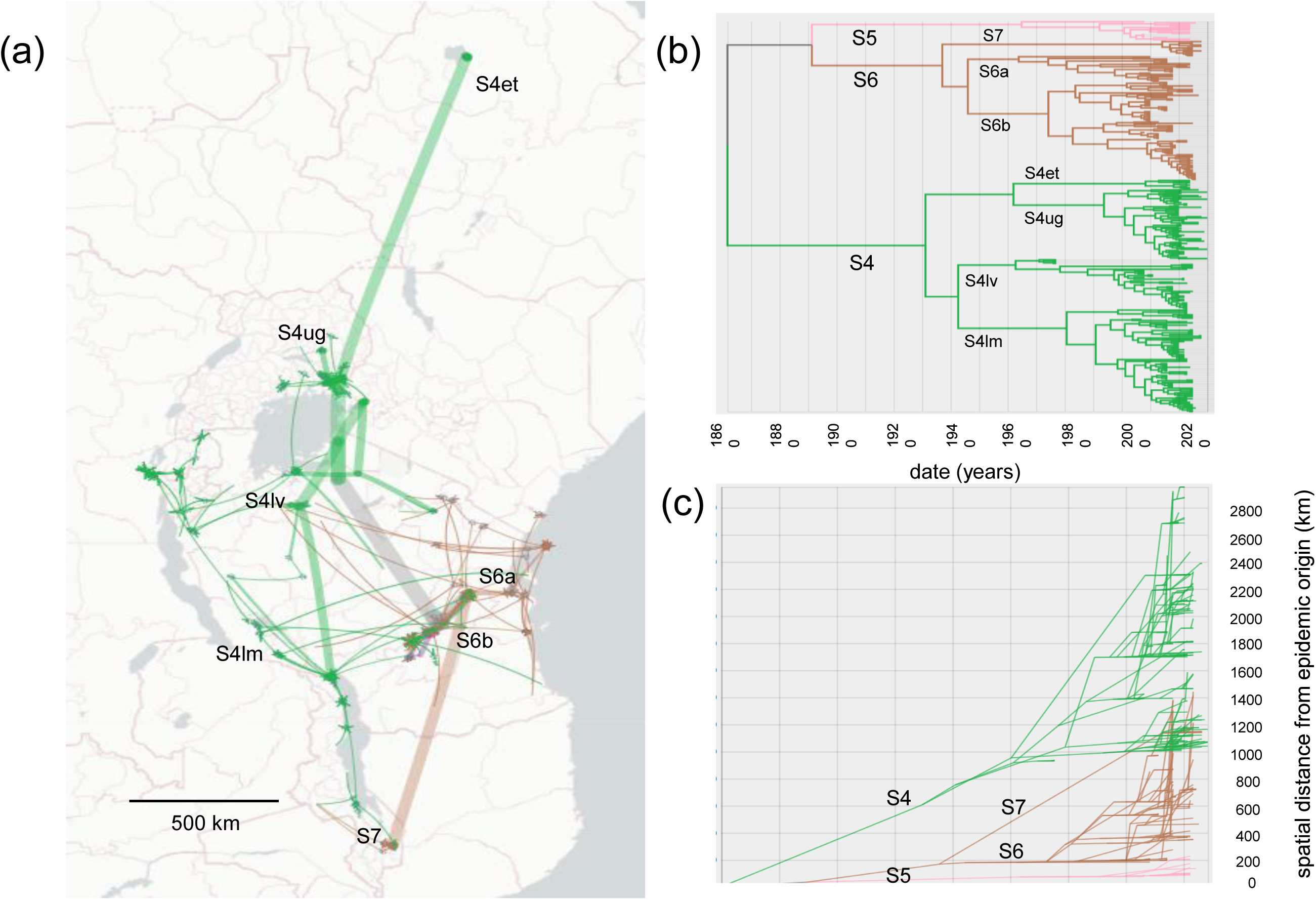
(a) Continuous phylogeographic reconstruction of the spatiotemporal dispersal of the three main lineages of RYMV in East Africa from 1850 to 2020 based on the coat protein gene sequences of 355 isolates collected from 1966 to 2019. The three lineages were distinguished by their colors: S4 (green), S6 (brown), S5 (pink). Note that the S5 lineage, confined to the Kilombero valley, is masked by the S6 lineage that has a broader distribution. The strains are indicated close to their emergence area (S4lv, S4lm, S4et, S4ug, S6a, S6b, S7). (b) Maximum credibility tree reconstructed from the dataset and used to generate the spatio-temporal dispersal (same color code for each lineage). (c) Migration curves over time (preorder transversal of the phylogenetic tree) for each parent/child node pair: X-axis = cumulative branch length (as dates in years), Y-axis= cumulative great-circle distances knowing latitudes and longitudes.

Additional sampling was performed from 2013 to 2017 at the Kilombero Rice Plantation (KPL) rice intensification station, located within the Kilombero Valley, which covers 5 000 ha of rice. A total of 75 isolates was collected in rice fields under intensifying schemes and in surrounding traditional rice fields. The coat protein gene of these isolates was sequenced. Almost all of the isolates belong to strain S6b already found in this region, one isolate of strain S6a, one S5 isolate and four S4lm isolates (data not shown). The absence of new strains and new variants indicate that the diversity of the virus in the Kilombero valley was adequately represented in the datasets used to reconstruct the phylogeography of RYMV in East Africa. The low proportion of isolates, other than of the S6b strain, shows that the circulation of strains in the Kilomobero rice valley was restricted, even in the present time.

### East African origin of the introduction of RYMV into West Africa and into Madagascar

The phylogeny of RYMV was reconstructed from the full-length sequences of 101 isolates representative of the genetic and geographic diversity of the virus in Africa: 50 isolates from East Africa, 45 from West Africa and six sequences from Madagascar. A major split in the phylogenetic network separates East Africa and Madagascar isolates from West Africa ones (Fig. 8a). The phylogeny is characterized by the nested position of the West African lineage within the East Africa lineage comprising the S4ug, S4mg and S4et strains (Fig. 8b). The S4mg strain in Madagascar resulted from a recombination between the S4ug strain and an unidentified minor parent 14]. The relationship between the West-African lineage and the S4et strain of this lineage persist after excluding the S4mg and S4ug sequences (Fig. 8c). This confirms the earlier assumptions, made on a restricted dataset, that RYMV in West Africa and in Madagascar originated in East Africa [14, 15], and brings information on the source lineage of RYMV in West Africa.

**Figure 8.**
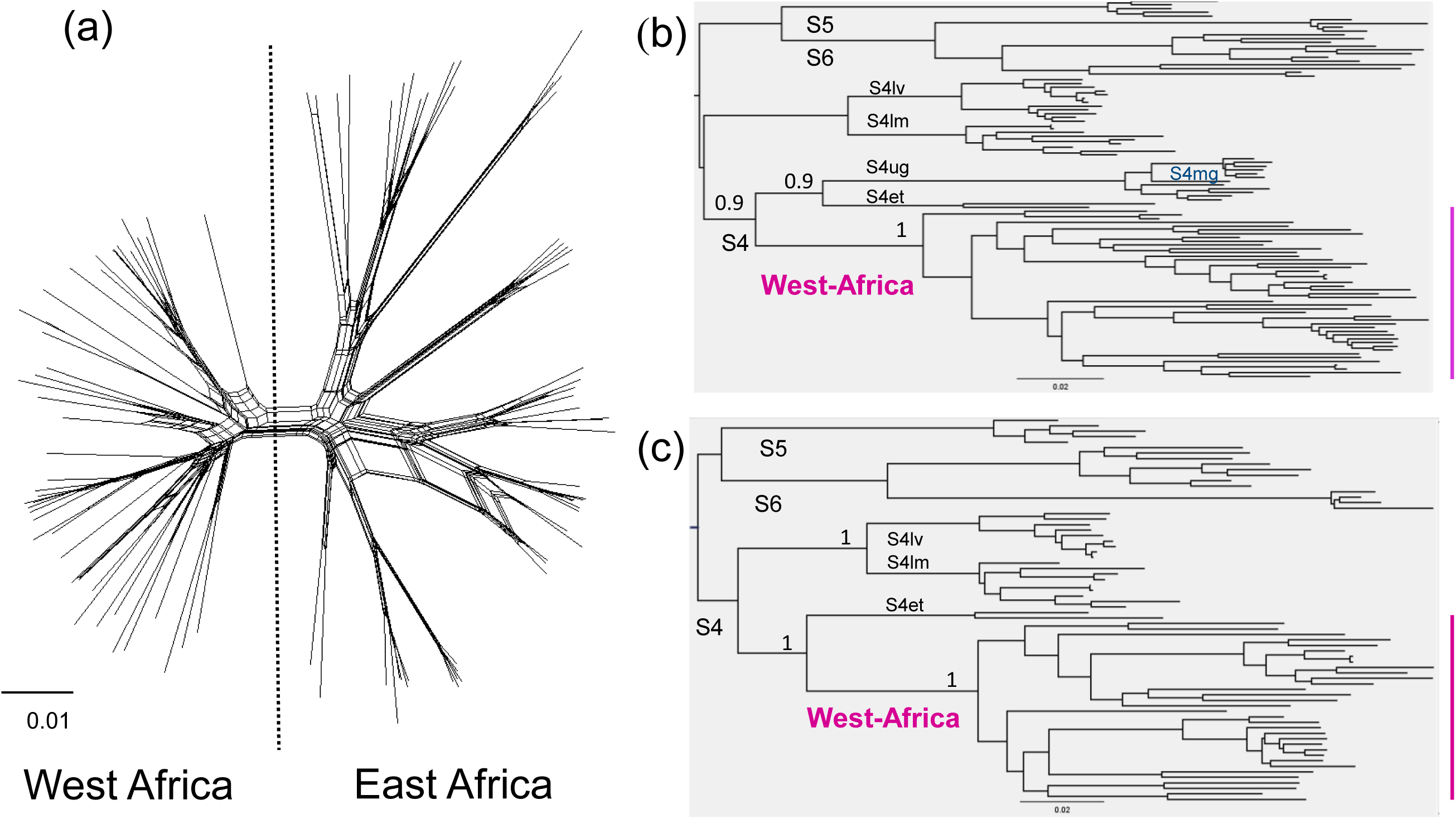
NeighborNet phylogenetic network of the 101 fully sequenced isolates of RYMV from East Africa and West Africa (purple) under a HKY85 distance model (8a). Maximum likelihood phylogeny tree reconstructed under a HKY85 model and mid-point rooting of the recombination-free full sequences obtained after removing the recombinant regions (8b), and after removing the recombinant sequences (8c).

We detailed the East African origin of the introduction of RYMV into West Africa and into Madagascar. Convergence of the analyses with the full dataset (AE335, WA335 and Mg94 dataset, i.e. a total of 784 coat protein gene sequences) was difficult to reach. So 50 isolates representative of the diversity of the virus in West Africa and 15 isolates representative of that in Madagascar were selected using PDA analyzer [37]. Then we reconstructed the phylogeography of the collection of the coat protein gene of 335 isolates from East Africa, the 50 representative isolates from West Africa and the 15 representative isolates from Madagascar, i.e. a total of 400 isolates. The phylogeographic model was that defined for the analysis of the EA335 sequence dataset (see Materials and Methods). The split between the East and the West African clades occurred in ca. 1868 (95% HPD 1831-1906) at the south of Lake Victoria (Fig. 9). The introduction of RYMV in the Niger Inner Delta in West Africa occurred in ca. 1892 (95% HPD, 1857-1909). The subsequent pattern of dispersion throughout West Africa and toward Central Africa mirrored the findings from comprehensive surveys conducted in West Africa [38, 39]. In Madagascar, RYMV was introduced in ca. 1980 (95% HPD 1975-1986) from the northeast of Lake Victoria. The introduction of RYMV in the northwest of Madagascar followed by the spread of the virus southward of the country is a scenario similar to that obtained from the comprehensive dataset [14].

**Figure 9.**
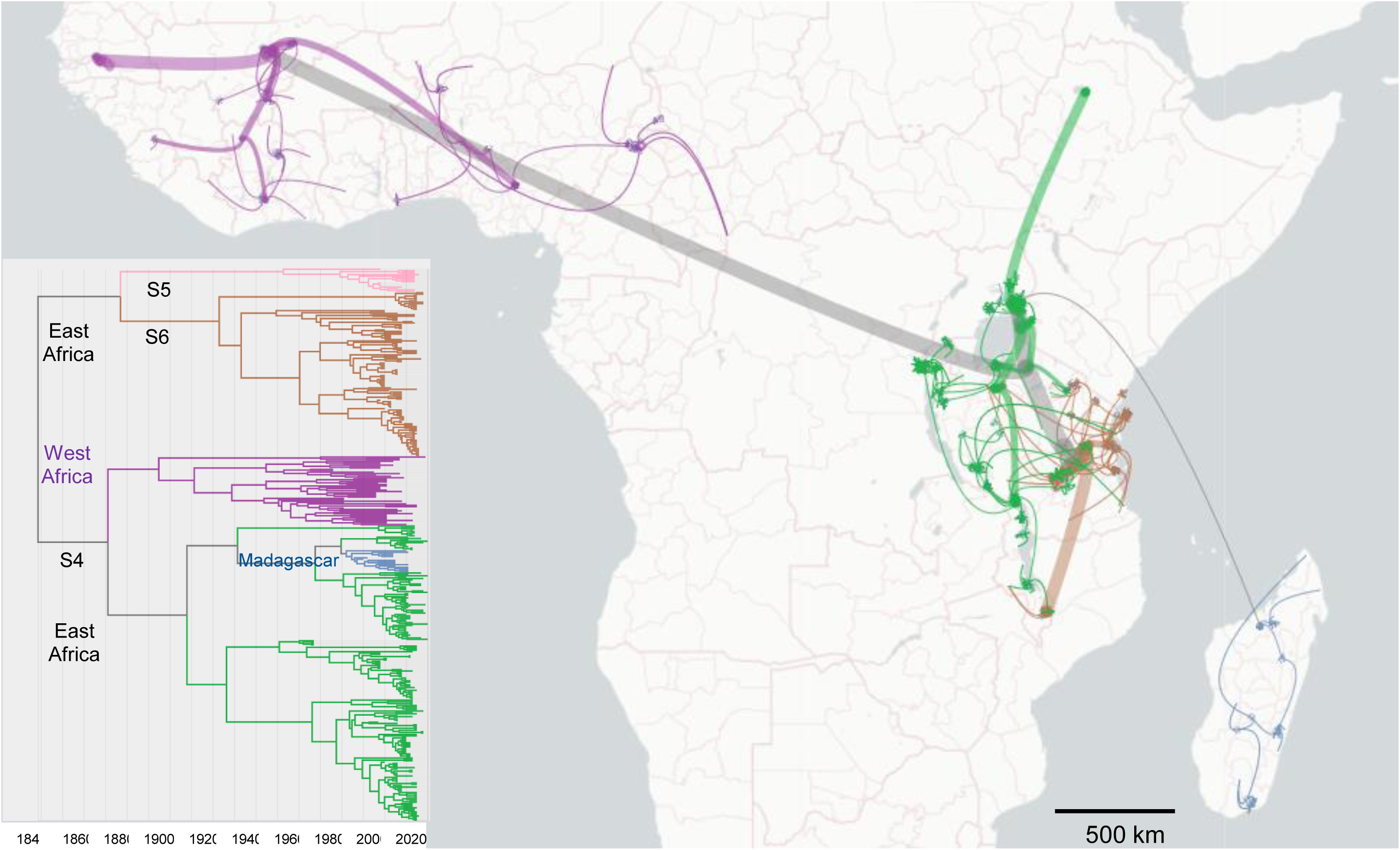
Dispersion of RYMV from East Africa (green, brown and pink for S4, S6 and S5 lineages, respectively) to West Africa (purple) and to Madagascar (blue) between 1850 and 2020 reconstructed from the ORF4 sequences of 335 isolates from East Africa, 50 isolates from West Africa and 15 isolates from Madagascar. The phylogeographic scenario are displayed with gradients linked to line thickness, curvature, opacity, and color. These gradients match the depth of the nodes in the phylogenetic tree, ranging from ancient to recent: path thickness ranges from wide to narrow, curvature varies from high to low, opacity shifts from light to strong, and color transitions from light to deep yellow. Inlet: Maximum credibility tree reconstructed from the dataset and used to generate the spatio-temporal dispersal (same lineage color code).

### Comparative dispersal statistics of RYMV

For RYMV, variation in isolation-by-distance (IBD) signals among the three regions (East Africa, West Africa and Madagascar) was apparent (from 0.35 to 0.56) (Table 2), yet it was less pronounced than that of rabies virus lineages in different host species (from 0.09 to 0.74) [33]. The IBD index was higher in West Africa (0.52 and 0.56 for the two datasets) than in East Africa (0.37) and in Madagascar (0.35) (Table 2). In East Africa, the recurrent long distance human-mediated dispersal of RYMV in a wide range of directions revealed by this study is consistent with a low IBD signal. The higher IBD signal estimated in West Africa may result from the dominant eastward long-distance transmission events establishing more local transmission chains [38, 39, 40]. The relatively low IBD signal estimated in Madagascar suggested that long-distance human-mediated dispersal, which blurred the relationship between geographical and genetic distances, is also frequent in a country where rice is widely grown. Diffusion coefficients in the three regions ranged from 1100 to 1600 km^2^ per year with overlapping 95% HPD intervals. This indicates a similar dispersal capacity of the virus across the three regions despite differences in IBD signal values, spatial scales, rice history and cultivation, and possibly also in dispersal modes.

**Table 2:**
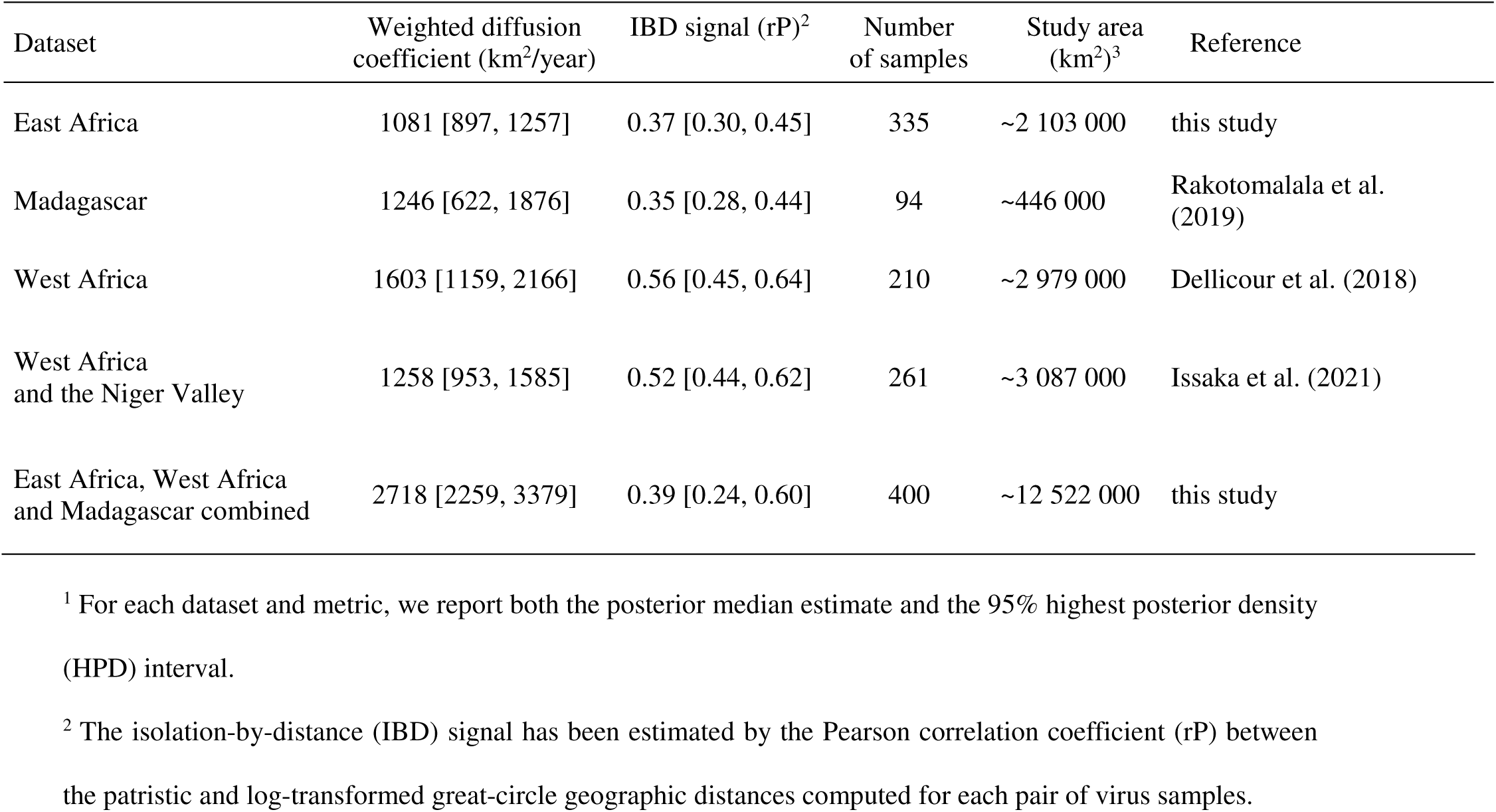
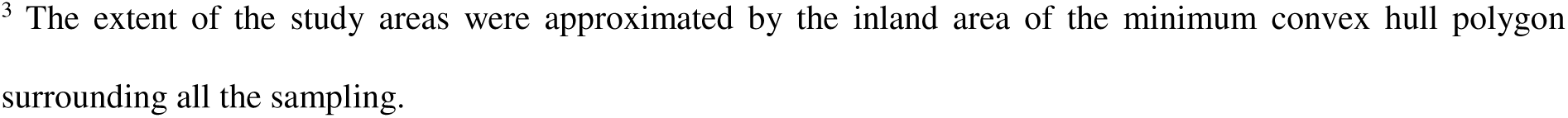
Dispersal metrics of RYMV in East Africa, West Africa, Madagascar and the three regions combined^1^.

| Dataset | Weighted diffusion coefficient (km<sup>2</sup>/year) | IBD signal (rP)<sup>2</sup> | Number of samples | Study area (km<sup>2</sup>)<sup>3</sup> | Reference |
| --- | --- | --- | --- | --- | --- |
| East Africa | 1081 [897, 1257] | 0.37 [0.30, 0.45] | 335 | ~2 103 000 | this study |
| Madagascar | 1246 [622, 1876] | 0.35 [0.28, 0.44] | 94 | ~446 000 | Rakotomalala et al. (2019) |
| West Africa | 1603 [1159, 2166] | 0.56 [0.45, 0.64] | 210 | ~2 979 000 | Dellicour et al. (2018) |
| West Africa and the Niger Valley | 1258 [953, 1585] | 0.52 [0.44, 0.62] | 261 | ~3 087 000 | Issaka et al. (2021) |
| East Africa, West Africa and Madagascar combined | 2718 [2259, 3379] | 0.39 [0.24, 0.60] | 400 | ~12 522 000 | this study |

<sup>1</sup> For each dataset and metric, we report both the posterior median estimate and the 95% highest posterior density (HPD) interval.
<sup>2</sup> The isolation-by-distance (IBD) signal has been estimated by the Pearson correlation coefficient (rP) between the patristic and log-transformed great-circle geographic distances computed for each pair of virus samples.
<sup>3</sup> The extent of the study areas were approximated by the inland area of the minimum convex hull polygon surrounding all the sampling.

The diffusion coefficient of RYMV for East Africa, West Africa and Madagascar combined (covering ca. 12 million km^2^) was 2700 km^2^/year, was twice that of each of three regions reflecting the inclusion of the long distance dispersal from East Africa towards West Africa and towards Madagascar (Table 2). The diffusion coefficient of RYMV was higher than that of several animal viruses including rabies in various hosts, even when occasional rapid long-distance dispersal events driven by human-mediated movements occurred [33]. Yet, the diffusion coefficient of RYMV was lower by an order of magnitude than that of animal viruses with long distance dispersal through animal trade or through migratory birds [33].

## Discussion

The seven periods identified in the phylogeographic reconstructions of RYVM in East Africa are interpreted as followed on the basis of the history of rice since the beginning of the 19^th^ century.

1. The postulated emergence of RYMV in the Eastern Arc Mountains [41] was confirmed and characterized in more details. The emergence of RYMV dates back to the mid-19th century. It took place near the Great Ruaha Escarpment of the Udzungwa Mountains adjacent to the Kilombero valley and close to the Morogoro region. Consistently, the insertion-deletion polymorphism originated in isolates of this region. The original host(s) of RYMV were located within the Eastern Afromontane biodiversity hotspot [15, 41]. Slash and burn rice cultivation practiced in the mid-1800s near the Great Ruaha Escarpment of the Udzungwa Mountains [28, 31, 42, 43] resulted in spillovers from the primary hosts of the virus, presumably of the Eragrostidae and related Poaceae families given RYMV host range, to cultivated rice. Primary hosts may also include the cultivated finger millet, a possible alternative host of RYMV, widespread in these mountains. This led to the introduction of RYMV in cultivated rice. The basal split between the S4, S5 and S6 lineages and the insertion-deletion polymorphism suggest (at least) three independent splillovers from the wild host(s) to cultivated rice. Strain specific poaceae species such as Dactyloctenium aegyptium have been identified [44]. Specific mutations of the S7 strain occurred at positions of the VPg under positive selection and experimentally associated to host range [45, 46]. Further studies are necessary to understand the genetic changes, if any, associated to host jumps and rice spillover.
2. Within a few decades, RYMV spread to the large adjacent Kilombero Valley where rice had been widely cultivated since the beginning of the 19^th^ century [28]. This is consistent with the movements of rice-growing populations between the Udzungwa Mountains and the Kilombero Valley [28].
3. Between the second half of the 19 ^th^ century and the beginning of the 20^th^ century, RYMV (lineage S4) spread to the south of Lake Victoria 400 km northward of the Kilombero Valley. Its dispersal followed the introduction of rice into the interior of the continent along the caravan routes linking the coastal strip to Lake Victoria [47, 48, 49]. There are indications that the caravan routes crossed the Kilombero valley [31]. Interestingly, the migration pathways of turnip mosaic virus (TuMV), a seed transmitted potyvirus, retraced some of the historical trade arteries of the Silk Road [50]. So, both RYMV and TuMV illustrate the link between plant virus dispersal and trade routes. During the same period, RYMV (lineage S6) was introduced in the Morogoro region 100 km eastward of the Kilombero Valley.
4. In the following decades, up to the middle of the 20^th^ century, RYMV dispersed at the east of Lake Victoria where rice had been introduced at the end of the 19^th^ century, probably by boat [51, 52]. Interestingly, the first field report of RYMV in East Africa was in 1966 near Kisumu also at the east of Lake Victoria [5].
5. At the beginning of the 20th century, RYMV was introduced from the Kilombero valley to the south of Lake Malawi. This movement is parallel and concomitant to the movement of military troops at the end of the First World War [53]. Feeding the troops was a major issue [54], rice was the main staple food for soldiers, and the Kilombero valley was the main rice production area. Accordingly, RYMV may have been dispersed through the transport of contaminated seeds during the First World War. An alternative hypothesis, linking virus dispersal to the Kilwa-Nyasa south caravan route, from Kiwa 300 km south of Zanzibar to the south of Lake Malawi (formerly named Lake Nyasa), is less likely. First, there is no firm indication that rice had been introduced at the south of Lake Malawi along this caravan route [55, 56]. Second, the peak of activity along the Kilwa-Nyasa south caravan route occurred in the second half of the 19^th^ century [55, 56] whereas the introduction of RYMV at the south of Lake Malawi occurred at the beginning of the 20^th^ century.
6. In the second half of the 20th century, RYMV was introduced into northern Ethiopia from the north of Lake Victoria. Large rice intensification schemes were established in the 1960s around Lake Victoria [5] and in the 1980s around Lake Tana in northern Ethiopia [57]. Seed exchanges between these intensification schemes were frequent and may have introduced RYMV in northern Ethiopia. The widespread cultivation of teff (Eragrostidae), a possible host of RYMV, in the Ethiopian highlands may have facilitated the dissemination.
7. RYMV then spread eastward and northward from the Morogoro region (strain S6a) and from the Kilombero Valley (strain S6b), and southward and westward from the Lake Victoria region (strains S4). RYMV finally became widespread throughout East Africa. This spread accompanied the expansion of rice cultivation of the last decades of the 20th century and of the beginning of the 21^st^ century. The slowdown in the dispersal rates of several strains of the S4 and S6 lineages over the past decades reflected a shift from an expansion phase to an endemic phase [32, 33] in RYMV epidemiology. This may be due to changes in the trade and market patterns [58] with a gradual change from long distance to local seed transport as rice became widely cultivated.

Altogether, the long-distance dispersal of RYMV was closely linked to rice movement in several regions, at different periods and under various circumstances. The dispersal of the RYMV was associated to rice introduction along the caravan routes from the Indian Ocean Coast to Lake Victoria in the second half of the 19^th^ century, to rice transport as troop staple food from the Kilombero valley towards the South of Lake Malawi at the end of the First World War, to seed exchanges between Lake Victoria and Ethiopia in the second half of the 20^th^ century. Alternative hypotheses involving beetle dissemination of the virus are unlikely given the paucity of rice cultivation and alternative hosts at these early periods between regions located several hundreds of kilometers apart. Seed transport could also account for short distance dispersal of the virus, in complement or as an alternative to beetle transmission, around Lake Victoria in the first half of the 20^th^ century and along the Indian Ocean in the past few decades. Finally, the ongoing transport of contaminated seeds could also explain the current generalized dispersal of RYMV throughout East Africa.

Early long-distance transport of contaminated seeds led to the dispersal of the main lineages in different directions and environments. So, the contrasting epidemic characteristics of the S4, S5 and S6 lineages resulted primarily from historical circumstances. In complement, the role of genetic properties of the strains should be explored. For instance, positive selection at positions of the VPg involved in the host range [45, 46] may have impacted the epidemiological characteristics of the S7 strain. The S5 lineage remained confined to the Kilombero valley, yet without becoming extinct, whereas the S4 lineage and later the S6 lineage were widely spread throughout East Africa. In West Africa, strain dominance was observed in competition experiments and by comparison of the virus titer of two strains [59]. Similar experiments with S5 and S6 and S4 isolates were not conclusive (Pinel-Galzi A, Fargette D, Hébrard E, unpublished results). Yet, these fitness tests may not encompass all the relevant traits involved in epidemiological differences and should be refined accordingly.

Phylogenetic analyses indicate that the West African RYMV diversity is nested as a monophyletic clade within the East African diversity, indicating an East African origin [15, 41]. The current phylogeographic study (i) confirms this hypothesis, (ii) brings information on the date, the location and the lineage of the origin, (iii) provides clues on the means of introduction, (iv) shows that the spread of RYMV throughout Africa can be reconstructed from a single RRW model. The origin of RYMV in East Africa was confirmed, based on a more representative dataset, by the nested phylogenetic position and the subset of insertion-deletion polymorphism of the West African clade. The phylogeography of RYMV throughout East Africa unveiled the role of rice transport in the long-distance movement of the virus. Similarly, the introduction of RYMV in West Africa may result from seed movements. Seed exchanges in Africa between agricultural stations, experimental farms, botanical and missionary gardens, even far apart, were frequent in the second half of the 19^th^ century [60, 61]. The Niger Inner Delta was a major destination for seeds from a wide range of origins, including Madagascar, because of rice intensification in this region since the end of the 19^th^ century [62, 63]. The origin of RYMV introduced in West Africa was located at the south of Victoria Lake at the end of the 19^th^ century, in Tanzania, which was under German occupation at that time. Contacts between German scientists and agricultural stations of the Niger Inner Delta at the end of the 19^th^ century are documented [61]. RYMV may have been introduced directly from this region, or indirectly through the Amani research station which was the main experimental station in East Africa [64]. The main trends of dispersal of RYMV throughout West Africa and Madagascar were captured within a single RRW model. This underlines the flexibility of the RRW models to reconstruct spatio-temporal dispersal when long-distance jumps, even transcontinental ones, occurred. The RRW that was used in this study is equivalent to a Cauchy process on a tree with rescaled branch lengths, which is a pure jump process allowing for long-distance dispersion events [65]. Our results also confirm the Ugandan origin of the Madagascar strain [14]. Increased seed exchanges between Madagascar and East African countries have occurred during the recent decades [66]. Transport of contaminated rice seeds may also account for this introduction.

RYMV dispersal through the transport of contaminated seeds shows the importance of this neglected virus transmission pathway in plant virus epidemiology [1]. This study also unveils the wide range of human activities, some of them unsuspected, involved in man-mediated movements of a plant virus. Consequently, a plant virus like RYMV has wide dispersal capacities, greater than or equal to that of most zoonotic viruses. Maize streak virus A (MSV-A) causes another major cereal disease in Africa [67]. It is transmitted by leafhoppers and not by seeds. Interestingly, introductions of MSV-A from Uganda to Madagascar and from Kenya to Ethiopia have been reported. These movements, parallels to those of RYMV, were also attributed to human-mediated introductions [66, 68]. It shows that the spread of RYMV and MSV-A, two major vector borne viruses, does not necessarily, or even primarily, depend on vector transmission.

An important pending question is why RYMV, given its wide dispersal capacities, is still restricted to Africa and does not have a worldwide distribution, unlike several sobemoviruses with similar biological properties. Long-distance movement of RYMV across the sea is possible but is quite a rare event as indicated by the unique successful introduction of the virus in West Africa. Rice was imported into America from West Africa in the 17^th^ century and early 18^th^ centuries, at least a century before the introduction of RYMV in West Africa. In the 19^th^ century, rice exports from East Africa were limited and directed towards regions where rice was not cultivated [30]. In the 20^th^ century, rice was imported into Africa rather than exported from this continent. Altogether, this explains why RYMV remained restricted to Africa. However, the risk of accidental introduction of RYMV into other continents subsists, in particular through the transfer of seeds for agronomical breeding which we have found to be a decisive means of long distance dissemination of the virus in Africa. Incidentally, sequences of MSV-A from the Philippines have recently been deposited on NCBI (PQ034624, OR983352 and eight others), a possible illustration of such human-mediated trans-continental introductions of vector borne plant viruses.

## Materials and Methods

### Datasets

A collection of 335 capsid protein gene sequences of East African isolates was built. The dataset consists of 240 sequences retrieved from NCBI (2024-06-01) supplemented by 95 sequences obtained in this study (Supplementary Table 1). The isolates originated from eight countries (Burundi, Republic of Congo, Ethiopia, Kenya, Malawi, Rwanda, Tanzania, Uganda). The sampling locations were distributed between latitudes 15.6° South and 11.9° North, and from longitudes 28.8° West to 39.8° East, ca. 3 000 kilometers in latitude x 1 200 km in longitude, i.e. a study area of ca. two million square kilometers. The isolates were sampled from 1966 to 2019 (54 years). A minor variant named S4ke recently identified in Kenya with a recombinant event in the capsid protein [69] was considered in this study (see below) but not included in this corpus.

A collection of full-length sequences of 101 isolates from Africa was set. This dataset consisted of sequences of 50 isolates of East Africa and the East of Central Africa (referred to as the East African dataset), 45 isolates of West Africa and the West of Central Africa (referred to as the West African dataset) and six sequences of Madagascar. Among the 50 full-length sequences of the East African dataset, 23 were sequenced for this study and 27 were retrieved from NCBI (Supplementary Table 2). This sample is representative of the genetic and geographic diversity of RYMV in East Africa, including the S4ke variant from western Uganda. No recombination signal was detected within the ORF2a and ORF2b genes.

In complement, a collection of 335 capsid protein gene sequences from West African isolates, was considered to estimate the substitution rate of RYMV (Supplementary Table 3). It consists of sequences retrieved from NCBI (2024-06-01), 261 sequences used in a study of the phylogeography in West Africa [38] complemented by 46 sequences of Ghana [39] and 28 sequences of Burkina Faso [40]. The 28 sequences from Burkina Faso were chosen from the 144 available on NCBI using the PDA software [37] in order to maximize the genetic diversity of the sample. In total, the samples were collected in 15 countries (Benin, Burkina Faso, Cameroon, Central African Republic, Chad, Gambia, Ghana, Guinea, Ivory Coast, Mali, Niger, Nigeria, Senegal, Sierra Leone, Togo). They were sampled between 1975 and 2021 (47 years).

Another estimate of the substitution rate was made from the collection of 94 capsid protein gene sequences of Malagasy isolates sampled between 1989 and 2017 (29 years) (Supplementary Table 4) that had been used to reconstruct the phylogeography of RYMV in Madagascar [14]. A temporal prior was introduced on the date of emergence of RYMV in Madagascar, the disease having been detected for the first time in the fields in the 1970s [14].

### Phylogenetic reconstructions

Sequences were aligned using CLUSTAL X with default parameters [70]. Maximum-likelihood (ML) phylogenetic trees were inferred using the PHYML-3.1 algorithm implemented in the SEAVIEW version 4.7 software [71] under a HKY85 substitution model. The ML trees were rooted at the point in the tree that minimizes the residual mean square of the root-to-tip distances. The neighborNet phylogenetic network of the 101 fully sequenced isolates was inferred [72] under an HKY85 distance model as implemented in SplitsTrees4. The neighborNet phylogenetic network illustrates the genetic relationships between sequences taking into account the conflicting phylogenetic signals that are possibly due to recombination events. The full-length sequence alignment was screened for recombination signals using RDP5 [73]. The default settings were used for each of the seven recombination detection algorithms that RDP incorporates, as was a Bonferroni corrected P-value cut-off of 0.05. Only recombination events detected by more than four of the seven methods implemented in RDP were considered. The phylogeny was reconstructed from recombinant-free alignments. This was obtained by removing either the recombinant sequences or the recombinant regions. In the latter case, it was checked that further recombination was excluded through the pairwise homoplasy (PHI) test [74] implemented in SplitsTrees4 [72]. Total, synonymous, and non-synonymous nucleotide diversity of ORF2a, ORF2b and ORF4 were assessed by DnaSP6 [75]. The ORF1, too short, too variable and with a low phylogenetic signal [76], was not considered.

### Bayesian evolutionary inferences

RYMV is a measurably evolving population, yet its overall temporal signal is weak [14, 15]. The temporal signal of each data set was assessed through tip cluster-randomization tests in BEAST [77] and by BETS [78] under a Bayesian statistical framework, and by TreeTime under a likelihood framework [79].

We subsequently reconstructed time-calibrated phylogenies using a Bayesian statistical framework implemented in BEAST version 1.10.4 software [80]. BEAST uses Markov Chain Monte Carlo integration to average over all plausible evolutionary histories for the data, as reflected by the posterior probability. All analyses were performed using the BEAGLE library to enhance computation speed [81, 82]. We specified an HKY85 substitution model with a discretized gamma distribution (four categories) to model rate heterogeneity across sites. To accommodate lineage-rate variation, an uncorrelated relaxed molecular clock that models the branch rate variation according to a lognormal distribution was specified [83]. The flexible nonparametric demographic skygrid prior was selected to accommodate for variation in the rate of coalescence through time [84]. Stationarity and mixing (e.g. based on effective sample sizes > 200 for the continuous parameters) were assessed using Tracer version 1.7 [85]. Consensus trees deriving from the phylogenies sampled during the MCMC analysis were obtained using TreeAnnotator version 1.10.4 [80].

To study the geographical spread of RYMV in continuous space in East Africa and to quantify its tempo and dispersal, we fitted a continuous phylogenetic diffusion model to the East African dataset, modelling the changes in coordinates (latitude and longitude) along each branch in the evolutionary history as a bivariate normal random deviate [22]. As a more realistic alternative to homogeneous Brownian motion, we adopted a "Cauchy" RRW extension that models variation in dispersal rates across branches by independently drawing branch-specific rate scalers of the RRW precision matrix from a gamma distribution with shape and scale equal to 1/2 to relax the assumption of a constant spatial diffusion rate throughout the whole tree [22]. Bayesian inference using continuous diffusion models yields a posterior distribution of the phylogenetic trees, each having ancestral nodes annotated with location estimates. The EA335 dataset was used to reconstruct the dispersal of RYMV throughout East Africa. A spatial jitter of 0.1° in latitude and in longitude (i.e. ca 10 km) was applied to the locations of the isolates. This degree of noise for identical coordinates was needed to avoid improper posteriors under the RRW model and associated inference problems [22, 86]. Evolaps2 was used to visualize the continuous phylogeographic reconstruction of the dispersal history of RYMV lineages [87, 88]. The phylogeography of RYMV in East Africa was reconstructed from the coat protein gene sequences of the full dataset (335 isolates), from the full genome and different ORFs of a subsample of the 50 isolates representative of the genetic and geographic diversity of the virus.

### Comparative dispersal statistics of RYMV

The phylogeography of RYMV has been reconstructed in East Africa (this study), in West Africa [38] and in Madagascar [14]. The dispersal capacity and dynamics in the three regions was compared through the weighted diffusion coefficient (km^2^/year) [15] and the isolation-by-distance (IBD) signal metric, two metrics identified as robust (to sampling intensity in particular) and measuring complementary aspects of the overall dispersal pattern [33]. The diffusion coefficient is an estimate of the dispersal capacity of the lineages. The IBD signal metric quantifies how the dispersal is spatially structured, i.e. the tendency of phylogenetically closely related tip nodes to be sampled from geographically close locations. An IBD signal was apparent through the relationships between geographic and genetic pairwise distances in a restricted dataset of sequences from East and West Africa [89]. The diffusion coefficient of RYMV virus over the three regions combined was also estimated. Then, the dispersal capacity of this plant virus was compared to that of a range of animal viruses in various host species [33].

## Acknowledgments

This work benefited from insightful discussions with Dick Peters over the years. Dick Peters passed away in November 2023.

This work was partly supported by the French National Research Agency as an “Investissements d’avenir” program (ANR-10-LABX-001-01 Labex Agro) coordinated by Agropolis Foundation (project no. 1504-004 E-SPACE) and by a bilateral project between Kenya and France (PHC PAMOJA 36128PK) cofunded by National Commission for Science, Technology and Innovation (NACOSTI) and Ministère de l’Europe et des Affaires Etrangères (MEAE). PR’s internship at the University of Montpellier was founded by the I-SITE MUSE through the Key Initiative “Data and Life Sciences”. SD acknowledges support from the Fonds National de la Recherche Scientifique (F.R.S.-FNRS, Belgium; grant n°F.4515.22), from the Research Foundation - Flanders (Fonds voor Wetenschappelijk Onderzoek - Vlaanderen, FWO, Belgium; grant n°G098321N), and from the European Union Horizon 2020 projects MOOD (grant agreement n°874850) and LEAPS (grant agreement n°101094685).

The funders had no role in the study design, data collection and interpretation, or the decision to submit the work for publication

## Supporting information

Supplementary Table 1. Name and accession number of the 335 isolates from East Africa.

Supplementary Table 2. Name and accession number of the 101 fully sequenced isolates from West Africa, East Africa and Madagascar.

Supplementary Table 3. Name and accession number of the 335 isolates from West Africa. Supplementary Table 4. Name and accession number of the 94 isolates of Madagascar.

Supplementary Table 5. Length, diversity and selection pressure of the ORF of Rice yellow mottle virus.

## References

1. Jones R. 2018. Plant and insect viruses in managed and natural environments: Novel and Neglected Transmission Pathways. Advances in Virus Research, 101: 149–187.

2. Peters D et al. 2024. The plant virus transmissions database. Journal of General Virology 105, doi 10.1099/jgv.0.001957

3. Sõmera M et al. 2021. ICTV Virus Taxonomy Profile: Solemoviridae. Journal of General Virology 102, 10.1099/jgv.0.001707

4. Abo M et al. 1997. Rice yellow mottle virus in Africa: evolution, distribution, economic significance and sustainable rice production and management strategies. Journal of Sustainable Agriculture 11, 85–111.

5. Bakker W. 1974. Characterization and ecological aspects of rice yellow mottle virus in Kenya. Agricultural Research Report 829, 1–152.

6. Allarangaye M et al. 2006. Evidence of non-transmission of rice yellow mottle virus through seeds of wild host species. Journal of Plant Pathology 88, 309–315.

7. Fauquet C and Thouvenel JC. 1977. Isolation of the rice yellow mottle virus in Ivory Coast. Plant Disease Reporter 61, 443–446.

8. Konaté G et al. 2001. Rice yellow mottle virus is seed-borne but not seed transmitted in rice seeds. European Journal of Plant Pathology 107, 361–364.

9. Sarra S and Peters D. 2003. Rice yellow mottle virus is transmitted by cows, donkeys, and grass rats in irrigated rice crops. Plant Disease, 87: 804–808.

10. Traoré O et al. 2009. A reassessment of the epidemiology of Rice yellow mottle virus following recent advances in field and molecular studies. Virus Research 141, 258– 267.

11. Cooper J and MacCallum F. 1984. Viruses and the Environment. Chapman and Hall. 182 pp.

12. Teakle D. 1986. Abiotic transmission of southern bean-mosaic virus in soil. Australian Journal of Biology Science 39, 353–360.

13. Traoré O et al. 2006. Rice seedbed as source of primary infection by Rice yellow mottle virus. European Journal of Plant Pathology 115, 181–186.

14. Rakotomalala M et al. 2019. Comparing patterns and scales of plant virus phylogeography: Rice yellow mottle virus in Madagascar and in continental Africa’. Virus Evolution 5, vez023.

15. Trovão N et al. 2015. Host ecology determines the dispersal patterns of a plant virus. Virus Evolution 1, vev016.

16. Abé Y. 2007. Le "Décorticage" du Riz. Editions de la Maison des Sciences de l’Homme. 588 pp.

17. Banwo O et al. 2004. Rice yellow mottle virus genus sobemovirus: a continental problem in Africa. Plant Protection Science 40, 26–36.

18. Richards P. 1986. Coping With Hunger. Hazard and Experiment in an African Rice-Farming System. Allen and Unwin. 176 pp.

19. Pedgley D. 1982. Windborne Pests and Diseases. Meteorology of Airborne Organisms. John Willey and Sons. 250 pp.

20. Thresh JM. 1983. The long-range dispersal of plant viruses by arthropod vectors. Phil. Trans. R. Soc. Lond. B302497–528. 10.1098/rstb.1983.0071

21. Lemey P et al. 2009. Bayesian phylogeography finds its roots. PLoS Computational Biology 5, e1000520.

22. Lemey P et al. 2010. Phylogeography takes a relaxed random walk in continuous space and time. Molecular Biology and Evolution, 27, 1877–1885.

23. Fargette D et al. 2008. Rice yellow mottle virus, an RNA plant virus, evolves as rapidly as most RNA animal viruses. Journal of Virology 82, 3584–3589.

24. Ghafari M et al. 2024. Revisiting the origins of the Sobemovirus genus: A case for ancient origins of plant viruses. PLoS Pathogens 10.1371/journal.ppat.1011911

25. Carpenter A. 1978. The history of rice in Africa. In Rice in Africa. Eds I. W. Buddenhagen and Persley G.J. Academic Press pp 3–10.

26. Gilbert E. 2016. Rice, civilization and the Swahili Towns: anti-commodity and anti-State?. In: Hazareesingh, S., Maat, H. (eds) Local Subversions of Colonial Cultures. Cambridge Imperial and Post-Colonial Studies Series. Palgrave Macmillan, London. 10.1057/9781137381101_9

27. Portères R. 1971. Riz. In Atlas des Cultures Vivrières. Bertin J, Hémardinquer JJ, Keul M. Randles W. Ecole Pratique des Hautes Etudes. Paris.

28. Jatzold and Baum. 1968. The Kilombero Valley. Characteristic features of the economic geography of a semihumid East African flood plain and its margins. Weltforum Verlag. 147 pp.

29. Alpers E. A. 1984. The Western Indian Ocean as a regional food network in the nineteenth century. International Congress on Indian Ocean studies, Perth, Western Australia, 5-12 December 1984. Proceedings Section A: Resources, Environment and Economic Development, 20 pp.

30. Alpers E. A. 2009. East Africa and the Indian Ocean. Markus Wiener Publishers, Princeton, 241 pp.

31. Iliffe J. 1979. A Modern History of Tanganyika. Cambridge University Press. 616 pp.

32. Bastide P, Pauline Rocu, Johannes Wirtz, Gabriel W Hassler, Francois Chevenet, Denis Fargette, Marc A Suchard, Simon Dellicour, Philippe Lemey, Stephane Guindon. 2024. Modeling the velocity of evolving lineages and predicting dispersal patterns. https://www.biorxiv.org/content/biorxiv/early/2024/06/06/2024.06.06.597755.full.pdf

33. Dellicour S. et al. 2024. How fast are viruses spreading in the wild? https://www.biorxiv.org/content/10.1101/2024.04.10.588821v2.full.pdf

34. Pybus OG, et al. 2012. Unifying the spatial epidemiology and molecular evolution of emerging epidemics. Proceedings of the National Academy of Sciences of the United States of America.109, 15066–15071.

35. Pybus O et al. 2007. Phylogenetic evidence for deleterious mutation load in RNA viruses and its contribution to viral evolution. Mol. Biol. Evol. 24, 845–852.

36. Fargette D et al. 2004. Inferring the evolutionary history of rice yellow mottle virus from genomic, phylogenetic, and phylogeographic studies. J Virol. 78, 3252–3261. doi: 10.1128/JVI.78.7.3252-3261

37. Chernomor O et al. 2015. Split diversity in constrained conservation prioritization using integer linear programming. Methods Ecol. Evol., 6, 83–91. DOI: 10.1111/2041-210X.12299

38. Issaka S et al. 2021. Rivers and landscape ecology of a plant virus, Rice yellow mottle virus along the Niger Valley. Virus Evolution. 7, veab072 10.1093/ve/veab072

39. Omiat E et al. 2023. Genetic diversity and epidemic histories of rice yellow mottle virus in Ghana. Viruses, 329 199106 10.1016/j.virusres.2023.199106

40. Billard E et al. 2023. Dynamics of the rice yellow mottle disease in western Burkina Faso: Epidemic monitoring, spatio-temporal variation of viral diversity and pathogenicity in a disease hotspot. Virus Evolution 9 vead049, 10.1093/ve/vead049

41. Pinel-Galzi A et al. 2015. The biogeography of viral emergence: rice yellow mottle virus as a case study. Current Opinion in Virology 10, 7–13.

42. Paul JL. 2003. Anthropologie Historique des Hautes Terres de Tanzanie Orientale. Karthala. 343 pp.

43. Raison JP. 1994. Tanzanie: l’Ujamaa et ses lendemains. In Les Afriques au Sud du Sahara. Belin-Reclus pp 372–388.

44. Allarangaye M et al. 2007. Host range of rice yellow mottle virus in Sudano-Sahelian savannahs. Pakistan Journal of Biological Sciences 9, 1414–1421.

45. Ndikumana I et al. 2017. Complete genome sequence of a new strain of rice yellow mottle virus from Malawi, characterized by a recombinant VPg Protein. Genome Announc. 5,: e01198–17. doi: 10.1128/genomeA.01198-17.

46. Poulicard N et al. 2012. Historical contingencies modulate the adaptability of Rice yellow mottle virus. PLoS Pathog 8, e1002482. doi: 10.1371/journal.ppat.1002482. Epub 2012.

47. Finch J et al. 2017. Ecosystem change in the South Pare Mountain bloc, Eastern Arc Mountains of Tanzania. The Holocene 27, 6 10.1177/0959683616675

48. Roberts A. 1970. Nyamwezi trade. In Pre-Colonial African Trade. Essays on Trade in Central and Eastern Africa before 1900. Eds Richard Gry and David Birmingham. Oxford University Press. pp 39–74.

49. Wynne-Jones S and Croucher S. 2007. The central caravan route of Tanzania: A preliminary archaeological reconnaissance. Nyame Akuma 67, 91–95.

50. Kawakubo S et al. 2021. Genomic analysis of the brassica pathogen turnip mosaic potyvirus reveals its spread along the former trade routes of the Silk Road. PNAS 118(12) e2021221118.

51. Leclercq J. 1913. Aux Sources du Nil par le Chemin de Fer de l’Ouganda. 302 pp.

52. Tosh J. 1970. The Northern Interlacustrine Region. In Pre-Colonial African Trade. Essays on Trade in Central and Eastern Africa before 1900. Eds Richard Gry and David Birmingham. Oxford University Press. pp 103–118.

53. Von Lettow-Vorbec, P. 1920. My Reminiscences of East Africa. Pantianos Classics.

54. Anderson R. 2016. Norforce: Major General Edward Northey and the Nyasaland and NorthEastern Rhodesia Frontier Force, January 1916 to June 1918. Scientia Militaria vol 44, no 1, 2016, pp 47–80. doi:10.5787/44-1-1162.

55. Alpers E. A. 1975. Ivory and Slaves in East Central Africa. Changing Patterns of International Trade to the Later Nineteenth Century. Heinemann. 296 pp.

56. Biginagwa T and Mapunda B. 2017. The Kilwa–Nyasa caravan route. The long-neglected trading corridor in southern Tanzania. In Wynnes-Jones S and LaViolette A « The Swahili World », chapter 47, pp 541–554.

57. Alemu D et al. 2018. A historical analysis of rice commercialisation in Ethiopia: the case of the Fogera plain. APRA Working Paper 18. Future Agricultures Consortium, Brighton.https://opendocs.ids.ac.uk/opendocs/handle/20.500.12413/14283

58. Meillassoux C (ed). 1971. The Development of Indigenous Trade and Markets in West Africa. Routledge. 444 pp.

59. N’Guessan P et al. 2000. Evidence of the presence of two serotypes of rice yellow mottle sobemovirus in Côte d’Ivoire. European Journal of Plant Pathology 106, 167– 178.

60. Bonneuil C. 1997. Mettre en ordre et discipliner les tropiques: les sciences du végétal dans l’Empire français, 1870-1940. Thèse de Doctorat à l’Université de Paris VII. 563 pp.

61. Tourte R. 2019. La Période Coloniale et les Grands Moments des Jardins d’Essais. 621 pp.

62. Viguier P. 1939. La Riziculture Indigène au Soudan Français. Paris: Larose 134 pp.

63. Portères R. 1950. Vieilles agricultures de l’Afrique Intertropicale. L’Agronomie Tropicale, vol 5, n°9-10 pp 489–507.

64. Blais H. 2023. L’Empire de la Nature: une Histoire des Jardins Botaniques Coloniaux (Fin XVIIIe siècle–années 1930). Champ Vallon Editions, 380 pp.

65. Bastide P and Didier G. 2023. The Cauchy process on phylogenies: a tractable model for pulsed evolution, Systematic Biology 72, 1296–1315, 10.1093/sysbio/syad053

66. Oyeniran K et al. 2024. Movement of the A-strain maize streak virus in and out of Madagascar, Virology, 10.1016/j.virol.2024.110222.

67. Shepherd D. 2010. Maize streak virus: an old and complex ‘emerging’ pathogen. Molecular Plant Pathology 11(1), 1–12 DOI: 10.1111/J.1364-3703.2009.00568

68. Ketsela D et al. 2022. Molecular identification and phylogenetic characterization of A-strain isolates of maize streak virus from western Ethiopia. Archives of Virology 167, 2753–2759.

69. Adego A et al. 2018. Full-length genome sequences of recombinant and nonrecombinant sympatric strains of rice yellow mottle virus from Western Kenya. Genome Announcements 6, 8 e01508–17

70. Thompson, et al. 1994. CLUSTAL W: Improving the sensitivity of progressive multiple sequence alignment through sequence weighting, position-specific gap penalties and weight matrix choice. Nucleic Acids Research 22, 4673–80.

71. Gouy M et al. 2010. SeaView version 4: a multiplatform graphical user interface for sequence alignment and phylogenetic tree building. Molecular Biology and Evolution 27, 221–224.

72. Bryant D and Moulton V. 2004. Neighbor-net: an agglomerative method for the construction of phylogenetic networks. Molecular Biology and Evolution, 21: 255– 265.

73. Martin D et al. 2020. RDP5: a computer program for analysing recombination in, and removing signals of recombination from, nucleotide sequence datasets. Virus Evolution 7, veaa087.

74. Bruen T et al. 2006. A simple robust statistical test for detecting the presence of recombination. Genetics, 172: 2665–2681.

75. Rozas J et al. 2017. DnaSP 6: DNA sequence polymorphism analysis of large data sets. Mol. Biol. Evol. 34, 3299–3302 doi:10.1093/molbev/msx248

76. Pinel-Galzi A et al. 2009. Recombination, selection and clock-like evolution of Rice yellow mottle virus. Virology 394, 164–172.

77. Murray G et al. 2016. The effect of genetic structure on molecular dating and tests for temporal signal. Methods in Ecology and Evolution 7, 80–89.

78. Duchêne, et al. 2020. Bayesian evaluation of temporal signal in measurably evolving populations. Molecular Biology and Evolution 37, 3363–3379.

79. Sagulenko P et al. 2018. TreeTime: maximum-likelihood phylodynamic analysis. Virus Evolution 4, vex042, 10.1093/ve/vex042

80. Suchard M et al 2018. Bayesian phylogenetic and phylodynamic data integration using BEAST 1.10, Virus Evolution 4, vey016.

81. Ayres D et al. 2012. BEAGLE: an application programming interface and high-performance computing library for statistical phylogenetics. Systematic Biology 61, 170–173.

82. Suchard M and Rambaut A. 2009.Many-core algorithms for statistical phylogenetics. Bioinformatics 25, 1370–1376.

83. Drummond A. et al. 2006. Relaxed phylogenetics and dating with confidence. PLoS Biology, 4: e88.

84. Gill M et al. 2013. Improving Bayesian population dynamics inference: a coalescent-based model for multiple loci. Molecular Biology and Evolution 30, 713–724.

85. Rambaut A et al. 2018. Posterior summarisation in Bayesian phylogenetics using Tracer 1.7. Systematic Biology 67, syy032.

86. Fernandez C and Steel M. 2000. Bayesian regression analysis with scale mixtures of normal. Econometric Theory 16, 80–101.

87. Chevenet, et al. 2021. EvoLaps: a web interface to visualize continuous phylogeographic reconstructions. BMC Bioinformatics 22, 463. 10.1186/s12859-021-04386-z

88. Chevenet, et al. 2024. EvoLaps 2: Advanced phylogeographic visualization. Virus Evolution 10, 1 vead078, 10.1093/ve/vead078

89. Abubakar Z et al. 2003. Phylogeography of rice yellow mottle virus in Africa. Journal of General Virology 84, 733–743.

