## Supplemental tables for "Grains, trade and war in the multimodal transmission of Rice yellow mottle virus: an historical and phylogeographical retrospective"

Supplementary Table 1. NCBI reference numbers of the coat protein gene sequences of 335 isolates representative of the genetic and geographic diversity of RYMV in East Africa (EA335 dataset).

| | |
|---|---|
| Bu1 | HE654712 |
| Bu10 | HE654716 |
| Bu113 | PQ243868* |
| Bu13 | HE654718 |
| Bu16 | HE654719 |
| Bu17 | HE654717 |
| Bu2 | HE654713 |
| Bu4 | HE654714 |
| Bu7 | HE654715 |
| Co177 | PQ243869* |
| Co209 | KC788208 |
| Co223 | KC788209 |
| Co229 | KC788210 |
| Co241 | PQ243870* |
| Co262 | PQ243871* |
| Et105 | MH917946 |
| Et15 | MH917947 |
| Et19 | MH917949 |
| Et2 | KM017554 |
| Et20 | MH917950 |
| Et21 | KM017557 |
| Et3 | KM017555 |
| Et5 | KM017556 |
| Ke1 | AJ511805 |
| Ke102 | MG599279 |
| Ke11 | FN432857 |
| Ke12 | FN432858 |
| Ke13 | FN432859 |
| Ke2 | AJ885128 |
| Ke3 | AM931178 |
| Ke323 | MG599279 |
| Ke325 | PQ243872* |
| Ke332 | PQ243873* |
| Ke343 | PQ243874* |
| Ke345 | MG599280 |
| Ke4 | PQ243875* |
| Mw1 | PQ243876* |
| Mw10 | MF989228 |
| Mw11 | PQ243966* |
| Mw12 | KP274897 |
| Mw13 | KP274897 |
| Mw14 | PQ243877* |
| Mw2 | KP274893 |
| Mw202 | PQ243878* |
| Mw203 | PQ243879* |
| Mw23 | PQ243880* |
| Mw24 | PQ243881* |
| Mw3 | KP274894 |
| Mw30 | PQ243882* |
| Mw301 | PQ243883* |
| Mw304 | PQ243884* |

| | |
|---|---|
| Mw306 | PQ243885* |
| Mw311 | PQ243886* |
| Mw318 | PQ243887* |
| Mw324 | PQ243888* |
| Mw4 | KP274895 |
| Mw46 | PQ243889* |
| Mw5 | KP274896 |
| Mw6 | PQ243890* |
| Mw66 | PQ243891* |
| Mw73 | PQ243892* |
| Mw74 | PQ243893* |
| Mw78 | PQ243894* |
| Mw84 | PQ243895* |
| Mw9 | MF989228 |
| Mw90 | PQ243896* |
| Mw95 | PQ243897* |
| Mw98 | PQ243898* |
| Rw1 | GQ470541 |
| Rw101 | HQ650135 |
| Rw102 | HQ650136 |
| Rw103 | HQ650137 |
| Rw104 | PQ256776* |
| Rw105 | PQ256777* |
| Rw110 | HE654711 |
| Rw15 | HE654704 |
| Rw181 | HE654703 |
| Rw19 | HE654706 |
| Rw2 | GQ470542 |
| Rw201 | HE654721 |
| Rw205 | HE654722 |
| Rw206 | HE654723 |
| Rw208 | HE654720 |
| Rw30 | HE654709 |
| Rw331 | PQ243899* |
| Tz1 | AJ279938 |
| Tz10 | AJ511799 |
| Tz1004 | FN432880 |
| Tz101 | AJ884693 |
| Tz1017 | FN432881 |
| Tz102 | AJ884705 |
| Tz1032 | FN432882 |
| Tz104 | AJ884690 |
| Tz1041 | FN432883 |
| Tz1044 | HE963829 |
| Tz1048 | FN432884 |
| Tz1051 | FN432885 |
| Tz1061 | FN432886 |
| Tz1066 | HE963828 |
| Tz1067 | HE963830 |
| Tz1069 | FN432887 |
| Tz1070 | FN432888 |
| Tz1072 | FN432889 |
| Tz1073 | FN432890 |
| Tz1074 | FN432891 |
| Tz1075 | FN432892 |

| | |
|---|---|
| Tz1077 | FN432893 |
| Tz1082 | FN432894 |
| Tz1086 | FN432895 |
| Tz109 | AJ884694 |
| Tz1094 | FN432896 |
| Tz1095 | HE963827 |
| Tz1099 | FN432897 |
| Tz11 | AJ511800 |
| Tz1101 | FN432898 |
| Tz1102 | FN432899 |
| Tz1103 | FN432900 |
| Tz1107 | FN432901 |
| Tz111 | AJ884714 |
| Tz1110 | FN432902 |
| Tz1112 | FN432903 |
| Tz1113 | FN432904 |
| Tz1114 | FN432905 |
| Tz1116 | FN432906 |
| Tz1118 | FN432907 |
| Tz112 | AJ884712 |
| Tz1120 | FN432908 |
| Tz1122 | FN432909 |
| Tz1125 | FN432910 |
| Tz1127 | FN432911 |
| Tz113 | AJ884711 |
| Tz1130 | FN432912 |
| Tz118 | AJ884707 |
| Tz12 | AJ511801 |
| Tz121 | AJ884704 |
| Tz122 | AJ884688 |
| Tz124 | AJ884702 |
| Tz125 | AJ884701 |
| Tz127 | AJ876793 |
| Tz128 | AJ884689 |
| Tz129 | AJ884713 |
| Tz13 | AJ885154 |
| Tz130 | AJ884699 |
| Tz14 | AJ885155 |
| Tz15 | AJ885156 |
| Tz150 | PQ243900* |
| Tz151 | PQ243901* |
| Tz152 | PQ243902* |
| Tz16 | AJ885157 |
| Tz17 | AJ885158 |
| Tz18 | AJ885159 |
| Tz19 | AJ885160 |
| Tz2 | AJ279939 |
| Tz20 | AJ885161 |
| Tz200 | AM931220 |
| Tz201 | AM931221 |
| Tz202 | AM883057 |
| Tz203 | AM931222 |
| Tz2032 | PQ243903* |
| Tz2034 | PQ243904* |
| Tz204 | AM931223 |

| | |
|---|---|
| Tz2044 | PQ243905* |
| Tz2046 | PQ243906* |
| Tz205 | AM931224 |
| Tz2050 | PQ243907* |
| Tz2055 | PQ243908* |
| Tz206 | AM931225 |
| Tz2067 | PQ243909* |
| Tz207 | AM883097 |
| Tz208 | AM931227 |
| Tz2085 | KM487714 |
| Tz209 | AM883058 |
| Tz21 | AJ885162 |
| Tz210 | AM931228 |
| Tz2100 | PQ243910* |
| Tz2105 | PQ243911* |
| Tz211 | AM931229 |
| Tz2116 | PQ243912* |
| Tz212 | AM931230 |
| Tz213 | AM931231 |
| Tz214 | AM931232 |
| Tz215 | AM931233 |
| Tz216 | AM931234 |
| Tz217 | AM931235 |
| Tz218 | AM931236 |
| Tz219 | AM931237 |
| Tz22 | AJ885163 |
| Tz220 | AM931238 |
| Tz221 | AM931239 |
| Tz222 | AM931240 |
| Tz223 | AM931241 |
| Tz225 | FN432873 |
| Tz226 | FN432874 |
| Tz229 | FN432877 |
| Tz23 | AJ885165 |
| Tz231 | FN432878 |
| Tz232 | FN432879 |
| Tz24 | AJ885165 |
| Tz3 | AJ279940 |
| Tz3001 | PQ243913* |
| Tz3004 | PQ243914* |
| Tz3005 | PQ243915* |
| Tz3007 | PQ243916* |
| Tz3017 | PQ243917* |
| Tz3018 | PQ243918* |
| Tz3026 | KM487715 |
| Tz3052 | PQ243919* |
| Tz3073 | PQ243920* |
| Tz308 | MF447484 |
| Tz319 | PQ256781* |
| Tz333 | MF447490 |
| Tz4 | AJ511793 |
| Tz401 | MF447484 |
| Tz402 | MF447485 |
| Tz4022 | PQ256778* |
| Tz4026 | PQ243921* |

| | |
|---|---|
| Tz403 | MF447486 |
| Tz4031 | MF447486 |
| Tz405 | MF447487 |
| Tz407 | MF447488 |
| Tz408 | MF447489 |
| Tz416 | MF447490 |
| Tz421 | MF447491 |
| Tz429 | MF447492 |
| Tz439 | PQ243922* |
| Tz441 | MF447493 |
| Tz445 | MF447494 |
| Tz449 | MF447495 |
| Tz450 | MF447496 |
| Tz452 | MF447497 |
| Tz454 | MF447498 |
| Tz460 | MF447499 |
| Tz461 | MF447500 |
| Tz463 | MF447501 |
| Tz483 | MF447502 |
| Tz486 | MF447503 |
| Tz5 | AJ511794 |
| Tz5001 | PQ243923* |
| Tz5004 | PQ243924* |
| Tz5007 | PQ243925* |
| Tz5008 | PQ243926* |
| Tz5009 | PQ243927* |
| Tz501 | PQ256779* |
| Tz5010 | PQ243928* |
| Tz5011 | PQ243929* |
| Tz5012 | PQ243930* |
| Tz5013 | PQ243931* |
| Tz503 | MF447504 |
| Tz504 | MF447505 |
| Tz505 | PQ243932* |
| Tz506 | PQ243933* |
| Tz507 | MF447506 |
| Tz510 | MF447508 |
| Tz512 | MF447509 |
| Tz513 | PQ243934* |
| Tz515 | MF447510 |
| Tz516 | MF447511 |
| Tz525 | MF447514 |
| Tz520 | MF447512 |
| Tz523 | MF447513 |
| Tz526 | MF447515 |
| Tz533 | MF447516 |
| Tz534 | PQ243935* |
| Tz539 | MF447517 |
| Tz543 | MF447518 |
| Tz554 | MF447519 |
| Tz561 | PQ243936* |
| Tz586 | PQ256780* |
| Tz6 | AJ511795 |
| Tz607 | MF447521 |
| Tz608 | MF447522 |

| | |
|---|---|
| Tz619 | MF447525 |
| Tz651 | MF447526 |
| Tz7 | AJ511796 |
| Tz701 | MF447527 |
| Tz702 | MF447528 |
| Tz8 | AJ511797 |
| Tz801 | MF447529 |
| Tz9 | AJ511798 |
| Ug1 | AM114523 |
| Ug1001 | PQ243937* |
| Ug1002 | PQ243938* |
| Ug1003 | PQ243939* |
| Ug101 | KM487718 |
| Ug102 | KM487718 |
| Ug103 | KM487719 |
| Ug104 | KM487720 |
| Ug105 | KM487721 |
| Ug107 | KM487738 |
| Ug108 | KM487722 |
| Ug109 | KM487723 |
| Ug110 | KM487724 |
| Ug142 | KM487725 |
| Ug143 | KM487726 |
| Ug144 | KM487727 |
| Ug145 | KM487728 |
| Ug146 | KM487729 |
| Ug148 | KM487730 |
| Ug18 | KM487711 |
| Ug2 | AM114524 |
| Ug201 | KM487731 |
| Ug204 | KM487732 |
| Ug205 | KM487733 |
| Ug207 | KM487712 |
| Ug210 | KM487734 |
| Ug211 | KM487735 |
| Ug212 | KM487736 |
| Ug220 | KM487737 |
| Ug230 | KM487713 |
| Ug317 | PQ243940* |
| Ug324 | PQ243941* |
| Ug332 | PQ243942* |
| Ug401 | AB981479 |
| Ug402 | AB985686 |
| Ug408 | AB980006 |
| Ug410 | AB980004 |
| Ug412 | AB980003 |
| Ug413 | AB980005 |
| Ug414 | AB980010 |
| Ug421 | AB980007 |
| Ug423 | PQ243943* |
| Ug425 | AB980008 |
| Ug426 | PQ243944* |
| Ug436 | PQ243945* |
| Ug439 | AB980009 |
| Ug440 | PQ243946* |

| | |
|---|---|
| Ug441 | PQ243947* |
| Ug442 | PQ243948* |
| Ug445 | PQ243949* |
| Ug460 | PQ243950* |
| Ug461 | PQ243951* |
| Ug463 | PQ243952* |
| Ug473 | PQ243953* |
| Ug484 | PQ243954* |
| Ug50 | KM487717 |

*this study

Supplementary Table 2. NCBI references of the full-length sequences of 101 isolates representative of the genetic and geographic diversity of RYMV in East Africa, West Africa and Madagascar.

| East Africa | | West Africa | | Madagascar | |
|---|---|---|---|---|---|
| Bu13 | PQ256769* | Be125 | MZ172961 | Mg1 | AJ608210 |
| Bu16 | PQ243960* | Be127 | MZ172956 | Mg2 | AJ608211 |
| Co177 | PQ243955* | Be136 | MZ172957 | Mg11 | JX966244 |
| Co223 | PQ243961* | BF_MP1401 | OQ858571 | Mg15 | JX966245 |
| Co229 | PQ243962* | BF_MP1493 | OQ858572 | Mg16 | AM883056 |
| Et20 | MH917950 | BF_MP3582 | OQ858573 | Mg20 | JX966246 |
| Et5 | MH917946 | BF1 | AM883059 | | |
| Ke101 | MG599277 | BF5 | MZ172956 | | |
| Ke105 | MG599278 | BF801 | MZ172959 | | |
| Ke12 | FN432839 | BF810 | MZ172958 | | |
| Ke2 | MG599276 | Ca5 | MZ172967 | | |
| Ke323 | MG599279 | Ce14 | MZ172968 | | |
| Ke345 | MG599280 | Ce26 | MZ172955 | | |
| Mw10 | MF989228 | CI4 | AJ608206 | | |
| Mw11 | PQ243966* | CI63 | AJ608207 | | |
| Mw306 | PQ256770* | CIa | AJ608219 | | |
| Mw324 | PQ279516* | CIb | L20893 | | |
| Mw9 | PQ243967* | Ga4 | FN432838 | | |
| Rw110 | PQ243963* | Gh113 | OQ225942 | | |
| Rw181 | PQ256771* | Gh124 | OQ225943 | | |
| Tz101 | MT701719 | Gh133 | OQ225944 | | |
| Tz11 | AJ608215 | Gh140 | OQ225945 | | |
| Tz1102 | JX966248 | Gh142 | OQ225946 | | |
| Tz1103 | PQ243956* | Ma10 | AJ608208 | | |
| Tz122 | OK181770 | Ma105 | MZ172969 | | |
| Tz125 | PP400330 | Ma203 | FN432840 | | |
| Tz127 | AJ876793 | Ma701 | MZ172970 | | |
| Tz16 | JX961551 | Ma77 | AJ608209 | | |
| Tz18 | AJ877020 | Ng101 | MZ172963 | | |
| Tz202 | AM883057 | Ng102 | *MZ172960* | | |
| Tz209 | AM883058 | Ng105 | MZ172964 | | |
| Tz21 | JX966247 | Ng106 | MF784437 | | |
| Tz211 | PQ279515* | Ng109 | MF784438 | | |
| Tz3 | AJ608216 | Ng119 | MZ172965 | | |
| Tz3001 | PQ256772* | Ng18 | FN432841 | | |
| Tz3005 | PQ256773* | Ng218 | MZ172954 | | |
| Tz3017 | PQ256774* | Ni1 | AJ608212 | | |
| Tz3018 | PQ256775* | Ni2 | AJ608213 | | |
| Tz319 | PQ256781* | Nia | U23142 | | |
| Tz421 | PQ243964* | Se1 | MN233654 | | |
| Tz429 | PQ243957* | Se5 | MN233655 | | |
| Tz454 | PQ243958* | SL4 | AJ608214 | | |
| Tz5 | AJ608217 | Tc28 | FN432837 | | |
| Tz508 | PQ243965* | Tg1 | MZ172966 | | |
| Tz8 | AJ608218 | Tg274 | MF784441 | | |
| Ug1 | KM487710 | | | | |
| Ug104 | PQ243959* | | | | |

| | |
|---|---|
| Ug148 | KM487711 |
| Ug207 | KM487712 |
| Ug23 | KM487713 |

* this study

Supplementary Table 3. NCBI reference numbers of the coat protein gene sequences of 335 isolates representative of the genetic and geographic diversity of RYMV in West Africa (WA335 dataset).

| | |
|---|---|
| Be1 | AJ885087 |
| Be120 | MZ172951 |
| Be123 | MZ172952 |
| Be125 | MZ172961 |
| Be127 | MZ172956 |
| Be136 | MZ172957 |
| Be2 | AJ885088 |
| Be27 | FN432845 |
| Be3 | AM931172 |
| Be4 | FN432842 |
| Be5 | FN432843 |
| Be6 | FN432844 |
| BF1 | AM883059 |
| BF2 | AJ885089 |
| BF5 | MZ172956 |
| BF570 | AJ885091 |
| BF572 | AJ885092 |
| BF682 | AJ885093 |
| BF702 | AM931174 |
| BF708 | OQ716830 |
| BF710 | OQ716832 |
| BF801 | MZ172959 |
| BF802 | FN432847 |
| BF803 | FN432848 |
| BF804 | FN432849 |
| BF810 | MZ172958 |
| BFef0333 | OQ716835 |
| BFef0738 | OQ716837 |
| BFmp0146 | OQ716848 |
| BFmp1397 | OQ716881 |
| BFmp1410 | OQ716868 |
| BFmp1451 | OQ716886 |
| BFmp1458 | OQ716887 |
| BFmp1824 | OQ716890 |
| BFmp1838 | OQ716895 |
| BFmp1878 | OQ716897 |
| BFmp1891 | OQ716900 |
| BFmp1892 | OQ716901 |
| BFmp2867 | OQ716903 |
| BFmp2899 | OQ716905 |
| BFmp2901 | OQ716907 |
| BFmp2952 | OQ716909 |
| BFmp2954 | OQ716910 |
| BFmp2966 | OQ716917 |
| BFmp2970 | OQ716918 |
| BFmp3584 | OQ716928 |
| BFmp4199 | OQ716932 |
| BFmp4200 | OQ716933 |
| BFmp4469 | OQ716949 |
| BFmp4477 | OQ716956 |
| BFmp4479 | OQ716958 |
| BFoq716904 | OQ716904 |
| Ca12 | MZ172914 |

| | |
|---|---|
| Ca2 | MZ172917 |
| Ca21 | MZ172915 |
| Ca22 | MZ172916 |
| Ca30 | AJ317950 |
| Ca38 | AJ306735 |
| Ca43 | MZ172918 |
| Ca5 | MZ172967 |
| Ca52 | MZ172919 |
| Ca54 | AJ317951 |
| Ce1 | KF054740 |
| Ce13 | MF066661 |
| Ce14 | MZ172968 |
| Ce163 | MF066693 |
| Ce17 | MF066664 |
| Ce170 | MF066694 |
| Ce175 | MF066695 |
| Ce177 | MF066697 |
| Ce181 | MF066698 |
| Ce183 | MF066699 |
| Ce19 | MF066665 |
| Ce2 | KF054741 |
| Ce20 | MF066666 |
| Ce22 | MF066667 |
| Ce226 | MF066669 |
| Ce228 | MF066671 |
| Ce26 | MZ172955 |
| Ce28 | MF066670 |
| Ce31 | KF054745 |
| Ce32 | KF054742 |
| Ce37 | MF066675 |
| Ce4 | KF054743 |
| Ce46 | MF066676 |
| Ce5 | MF066660 |
| Ce61 | MF066681 |
| Ce63 | MF066682 |
| Ce64 | MF066683 |
| Ce66 | MZ172953 |
| Ce72 | MF066684 |
| Ce93 | MF066689 |
| CI1* | AJ279902 |
| CI10 | AJ279911 |
| CI101 | AJ885099 |
| CI104 | AJ885100 |
| CI105 | AJ885101 |
| CI106 | AJ885102 |
| CI109 | AJ885103 |
| CI11 | AJ279912 |
| CI110 | AJ885104 |
| CI111 | AJ885105 |
| CI112 | AJ885106 |
| CI113 | AJ885107 |
| CI114 | AJ885108 |
| CI115 | AJ885109 |
| CI116 | AJ885110 |
| CI117 | AJ885111 |
| CI118 | AJ885112 |
| CI12 | AJ279913 |

| | |
|---|---|
| CI121 | AJ885113 |
| CI129 | AJ885114 |
| CI13 | AJ279914 |
| CI138 | AJ885115 |
| CI139 | AJ885116 |
| CI14* | AJ279915 |
| CI15* | AJ279916 |
| CI151 | AJ885117 |
| CI152 | AJ885118 |
| CI153 | AJ885119 |
| CI154 | AJ885120 |
| CI155 | AJ885121 |
| CI156 | AJ885122 |
| CI157 | AJ885123 |
| CI16 | AJ279917 |
| CI17 | AJ279918 |
| CI2 | AJ279903 |
| CI3 | AJ279904 |
| CI4 | AJ608206 |
| CI46 | AJ885094 |
| CI47 | AJ885095 |
| CI5 | AJ279906 |
| CI6 | AJ279907 |
| CI63 | AJ608207 |
| CI65 | AJ885096 |
| CI66 | AJ885097 |
| CI67 | AJ885098 |
| CI68 | AM931175 |
| CI7 | AJ279908 |
| CI8 | AJ279909 |
| CI9 | AJ279910 |
| CIa | AJ608219 |
| CIb | L20893 |
| Ga1 | AM765810 |
| Ga2 | AM765811 |
| Ga3 | AM765812 |
| Ga4 | FN432838 |
| Gh1 | AJ279919 |
| Gh101 | OQ200628 |
| Gh102 | OQ200629 |
| Gh103 | OQ200630 |
| Gh105 | OQ200631 |
| Gh106 | OQ200632 |
| Gh109 | OQ200633 |
| Gh111 | OQ200634 |
| Gh112 | OQ200635 |
| Gh113 | OQ200636 |
| Gh115 | OQ200637 |
| Gh116 | OQ200638 |
| Gh117 | OQ200639 |
| Gh118 | OQ200640 |
| Gh120 | OQ200641 |
| Gh121 | OQ200642 |
| Gh122 | OQ200643 |
| Gh124 | OQ200644 |
| Gh125 | OQ200645 |
| Gh126 | OQ200646 |

| | |
|---|---|
| Gh127 | OQ200647 |
| Gh128 | OQ200648 |
| Gh129 | OQ200649 |
| Gh133 | OQ200650 |
| Gh138 | OQ200651 |
| Gh139 | OQ200652 |
| Gh140 | OQ200653 |
| Gh141 | OQ200654 |
| Gh142 | OQ200655 |
| Gh3227 | OQ200620 |
| Gh3229 | OQ200621 |
| Gh3244 | OQ200622 |
| Gh3245 | OQ200623 |
| Gh3387 | OQ200624 |
| Gh3397 | OQ200625 |
| Gh3399 | OQ200626 |
| Gh3416 | OQ200627 |
| Gh46 | OQ200610 |
| Gh47 | OQ200611 |
| Gh48 | OQ200612 |
| Gh49 | OQ200613 |
| Gh50 | OQ200614 |
| Gh51 | OQ200615 |
| Gh52 | OQ200616 |
| Gh53 | OQ200617 |
| Gh54 | OQ200618 |
| Gh55 | OQ200619 |
| Gu1 | AJ279920 |
| Gu101 | FN432854 |
| Gu107 | FN432855 |
| Gu111 | FN432856 |
| Gu201 | FN432857 |
| Gu21 | AJ885124 |
| Gu26 | AJ885125 |
| Gu28 | AJ885126 |
| Gu29 | AJ885127 |
| Gu30 | AM931176 |
| Gu31 | AM931177 |
| Gu32 | AM931178 |
| Ma1 | AJ279926 |
| Ma10 | AJ608208 |
| Ma105 | MZ172969 |
| Ma144 | AJ885136 |
| Ma145 | AJ885137 |
| Ma146 | AJ885138 |
| Ma148 | AJ885139 |
| Ma149 | AJ885140 |
| Ma171 | AJ885141 |
| Ma175 | AJ885142 |
| Ma179 | AJ885143 |
| Ma180 | AJ885144 |
| Ma185 | AJ885145 |
| Ma2 | AJ279927 |
| Ma201 | AM931178 |
| Ma202 | AM931180 |
| Ma203 | FN432840 |
| Ma204 | AM931182 |

| | |
|---|---|
| Ma205 | AM931183 |
| Ma206 | AM931184 |
| Ma207 | AM931185 |
| Ma208 | AM931186 |
| Ma209 | AM931187 |
| Ma210 | AJ885146 |
| Ma25 | MZ172920 |
| Ma274 | MZ172921 |
| Ma29 | AJ885132 |
| Ma3 | AJ279928 |
| Ma301 | FN432860 |
| Ma302 | FN432861 |
| Ma303 | FN432862 |
| Ma304 | FN432863 |
| Ma305 | FN432864 |
| Ma34 | MZ172922 |
| Ma389 | MZ172923 |
| Ma4 | AJ279929 |
| Ma403 | MZ172924 |
| Ma41 | AJ885133 |
| Ma421 | MZ172925 |
| Ma43 | MZ172926 |
| Ma45 | MZ172927 |
| Ma479 | MZ172928 |
| Ma5 | AJ279930 |
| Ma6 | AJ279931 |
| Ma7 | AJ885129 |
| Ma701 | MZ172970 |
| Ma77 | AJ608209 |
| Ma8 | AJ885130 |
| Ma9 | AJ885131 |
| Ma98 | AJ885134 |
| Ng10 | AM931218 |
| Ng101 | MZ172963 |
| Ng102 | MZ172960 |
| Ng103 | MZ172931 |
| Ng104 | MZ172932 |
| Ng105 | MZ172964 |
| Ng106 | MF784437 |
| Ng109 | MF784438 |
| Ng11 | AM931219 |
| Ng110 | MZ172933 |
| Ng1101 | MZ172939 |
| Ng111 | MZ172934 |
| Ng113 | MF461282 |
| Ng114 | MZ172935 |
| Ng115 | MZ172936 |
| Ng119 | MZ172965 |
| Ng18 | FN432841 |
| Ng20 | FN432865 |
| Ng201 | MZ172940 |
| Ng21 | FN432866 |
| Ng215 | MZ172941 |
| Ng217 | MZ172943 |
| Ng218 | MZ172954 |
| Ng22 | FN432867 |
| Ng221 | MZ172944 |

| | |
|---|---|
| Ng23 | FN432868 |
| Ng24 | FN432869 |
| Ng25 | MZ172929 |
| Ng250 | MZ172942 |
| Ng27 | FN432870 |
| Ng28 | FN432871 |
| Ng29 | FN432872 |
| Ng3 | AJ885147 |
| Ng30 | MZ172930 |
| Ng321 | MZ172937 |
| Ng335 | MZ172938 |
| Ng4 | AJ885148 |
| Ng6 | AJ885149 |
| Ng7 | AM931215 |
| Ng8 | AM931216 |
| Ng9 | AM931217 |
| Ni1 | AJ608212 |
| Ni106 | MZ172945 |
| Ni107 | MZ172946 |
| Ni108 | MZ172947 |
| Ni111 | MZ172948 |
| Ni118 | MZ172949 |
| Ni124 | MZ172950 |
| Ni131 | AJ884691 |
| Ni2 | AJ608213 |
| Ni4 | AJ885150 |
| Ni6 | AJ885151 |
| Nia | U23142# |
| Se1 | MN233654 |
| Se5 | MN233655 |
| Se8 | MH699983 |
| SL1 | AJ279935 |
| SL2 | AJ279936 |
| SL3 | AJ279937 |
| SL4 | AJ608214 |
| SL5 | AJ885152 |
| SL6 | AJ279934 |
| SL7 | AJ885153 |
| Tc11 | AJ317952 |
| Tc17 | AJ317953 |
| Tc24 | AJ317954 |
| Tc28 | FN432837 |
| Tc31 | FN432850 |
| Tc32 | FN432851 |
| Tc33 | FN432852 |
| Tc34 | FN432853 |
| Tg1 | MZ172966 |
| Tg12 | AJ885170 |
| Tg21 | AJ885172 |
| Tg247 | MF784439 |
| Tg274 | MF784441 |
| Tg4 | AJ885167 |
| Tg7 | AJ885168 |
| Tg9 | AJ885169 |

Supplementary Table 3. NCBI reference numbers of the coat protein gene sequences of 94 isolates representative of the genetic and geographic diversity of RYMV in Madagascar (Mg94 dataset).

| | |
|---|---|
| Mg1 | AJ279921 |
| Mg10 | AM931190 |
| Mg105 | JX961583 |
| Mg108 | MK942093 |
| Mg11 | AM931191 |
| Mg119 | JX961584 |
| Mg12 | AM931192 |
| Mg123 | JX961585 |
| Mg124 | JX961586 |
| Mg125 | JX961587 |
| Mg126 | JX961588 |
| Mg127 | JX961589 |
| Mg128 | JX961590 |
| Mg13 | AM931192 |
| Mg14 | AM931194 |
| Mg147 | JX961591 |
| Mg15 | AM931195 |
| Mg150 | JX961592 |
| Mg155 | JX961593 |
| Mg16 | AM883056 |
| Mg163 | JX961594 |
| Mg165 | JX961595 |
| Mg17 | AM931196 |
| Mg18 | AM931197 |
| Mg19 | AM931198 |
| Mg2 | AJ279922 |
| Mg20 | AM931199 |
| Mg201 | JX961596 |
| Mg202 | JX961597 |
| Mg21 | AM931200 |
| Mg210 | JX961598 |
| Mg22 | AM931201 |
| Mg23 | AM931202 |
| Mg24 | AM931203 |
| Mg25 | AM931204 |
| Mg26 | AM931205 |
| Mg27 | AM931206 |
| Mg28 | AM931207 |
| Mg29 | AM931208 |
| Mg3 | AJ279923 |
| Mg30 | AM931209 |
| Mg301 | MK942094 |
| Mg302 | MK942095 |
| Mg303 | MK942096 |
| Mg304 | MK942097 |
| Mg31 | AM93210 |
| Mg32 | AM93211 |
| Mg33 | AM93212 |
| Mg34 | AM93213 |
| Mg341 | MK942098 |
| Mg342 | MK942099 |
| Mg343 | MK942100 |
| Mg344 | MK942101 |

| | |
|---|---|
| Mg35 | AM93214 |
| Mg4 | AJ279924 |
| Mg40 | JX912946 |
| Mg401 | MK942104 |
| Mg402 | MK942102 |
| Mg403 | MK942103 |
| Mg41 | JX961552 |
| Mg410 | MK942106 |
| Mg419 | MK942105 |
| Mg42 | JX961553 |
| Mg43 | JX961554 |
| Mg44 | JX961555 |
| Mg45 | JX961556 |
| Mg46 | JX961557 |
| Mg47 | JX961558 |
| Mg48 | JX961559 |
| Mg49 | JX961560 |
| Mg5* | AJ279925 |
| Mg50 | JX961561 |
| Mg51 | JX961562 |
| Mg52 | JX961563 |
| Mg53 | JX961564 |
| Mg54 | JX961565 |
| Mg55 | JX961566 |
| Mg56 | JX961567 |
| Mg57 | JX961568 |
| Mg58 | JX961569 |
| Mg59 | JX961570 |
| Mg6 | AM931189 |
| Mg60 | JX961571 |
| Mg61 | JX961572 |
| Mg62 | JX961573 |
| Mg63 | JX961574 |
| Mg64 | JX961575 |
| Mg65 | JX961576 |
| Mg66 | JX961577 |
| Mg67 | JX961578 |
| Mg68 | JX961579 |
| Mg69 | JX961580 |
| Mg70 | JX961581 |
| Mg71 | JX961582 |

Supplementary Table 2. Length, diversity and selection pressure of the ORF of Rice yellow mottle virus[1]

| ORF | ORF2a | ORF2b | ORF4 |
|---|---|---|---|
| Nucleotide positions[2] | 610-2427 | 1986-3605 | 3448-4167 |
| Number of codons | 605 | 540 | 240 |
| Nucleotide diversity | 0.053 | 0.057 | 0.084 |
| Synonymous diversity ($d_s$) | 0.153 | 0.183 | 0.229 |
| Non-synonymous diversity ($d_n$) | 0.018 | 0.017 | 0.034 |
| Ratio $d_n/d_s$ | 0.12 | 0.09 | 0.15 |

[1]Estimated from sequences of 50 isolates representative of the genetic and geographic diversity of RYMV in East Africa.
[2]The nucleotide positions are those of the NCBI accession number PP400330.
